# Repeated climate-driven dispersal and speciation in peripheral populations of Pleistocene mastodons

**DOI:** 10.1101/2024.12.03.626650

**Authors:** Emil Karpinski, Sina Baleka, Andrew R. Boehm, Tim Fedak, Chris Widga, Hendrik N. Poinar

**Affiliations:** Department of Genetics, Harvard Medical School, Boston, MA, 02115, USA; McMaster Ancient DNA Centre, Departments of Biochemistry and Anthropology, McMaster University, Hamilton ON, L8S 4L9, Canada; Museum of Natural and Cultural History, University of Oregon, Eugene, OR, 97403, USA; Nova Scotia Museum of Natural History, Halifax, NS, BH3 3A6, Canada; Earth and Mineral Sciences Museum & Art Gallery, Penn State University, University Park, PA, 16802, USA; Michael G. DeGroote Institute for Infectious Disease Research, McMaster University, Hamilton ON, L8S 4L9, Canada

**Keywords:** ancient DNA, *Mammut americanum*, *Mammut pacificus*, Pleistocene, Phylogeography

## Abstract

Recent ancient DNA work has shed some light on the responses of mastodons to Pleistocene glacial/interglacial cycling but focused primarily on their expansion into Beringia. However, genetics has complicated our understanding of the relationships within *Mammut*, specifically between Pacific and American mastodon phylogeography and questioned whether these are in fact two separate species or regionally localized morphotypes. Here we expand on both avenues by sequencing and contextualizing the mitochondrial genome of a Pacific mastodon, as well as from several North American eastern specimens throughout the last 800 thousand years. We show that Pacific mastodons fall within a previously established, and deeply divergent mitochondrial clade, extending the range of this species into western Canada and potentially Mexico. We also present evidence for at least three discrete expansion events into northeastern coastal regions (i.e. Nova Scotia and the eastern continental shelf), and identify two new mastodon clades, which contain temporally distinct, but geographically co-occurrent specimens. This work sheds further light on mastodon taxonomy and phylogeography across North America throughout the Pleistocene, highlighting interglacial range expansion into northeastern America mirroring the effects on the western side of the continent (Beringia).

## 1. Introduction

Our understanding of the evolutionary relationships between extinct taxa and their extant relatives has undergone successive revisions as new paleontological and biomolecular data become available (see for example refs ^1,2^). Our increasing ability to recover and analyze ancient, tiny, degraded DNA fragments from hundreds of thousands of year-old remains has allowed us to answer previously unresolved questions, adding a significant complementary line of inquiry into palaeontological studies, especially when recovered remains are often few in number or too fragmentary for clear morphological identification. These molecular data also allow us to address questions about the relationships between and estimate divergence times of particular populations, study their demographics, as well as their responses to various ecological and anthropogenic pressures ^3–5^.

Our understanding of North American mastodon taxonomy has undergone substantial revisions. In the 19^th^ and early 20^th^ centuries, *Mammut* consisted of many species, which were subsequently synonymized into a single late Pleistocene species (*Mammut americanum*), before being split again into two co-occurring late Pleistocene taxa, the American mastodon (*Mammut americanum*) and the Pacific mastodon (*Mammut pacificus*) ^6–8^. Previous ancient DNA work on mastodon remains from the American Falls Reservoir, has suggested that these two taxa might not be separate species and instead represent discrete morphotypes ^9^. Instead, it did reveal that Idaho Falls mastodons were part of a predominantly marine isotope stage 5 (MIS 5) clade of mastodons that likely expanded from the contiguous USA, up along the Rockies, and into the Arctic in response to interglacial warming ^4,9^. However, previous work was stymied by recovery of genetic material only from very fragmentary remains which precluded clear morphological identification ^9^.

These studies, highlighting morphological and genetic variability in late Pleistocene mastodons, have suggested a much more complex evolutionary history than previously appreciated. At this time, it is unclear if and how much of the genetic diversity observed reflects species-level versus population-level divergence. Our study begins to disentangle these differences by focusing primarily on mastodons from the peripheries of their core geographic ranges (*M. americanum*: Great Lakes ^10^; *M. pacificus*: California and Idaho ^7^) to further our understanding of mastodon phylogeography. Here we present sequence data from a well-identified Pacific mastodon from Tualatin, Oregon and five additional American mastodons from Nova Scotia and the eastern seaboard, within which we have identified novel and deep patterns of expansion/extirpation in response to glacial cycling. In tandem, these data fill holes in our understanding of mastodon evolutionary relationships and highlight the responsiveness of these species to glacial/interglacial cycles at the coastal margins of the continent.

## 2. Methods

### 2.1 Sample acquisition and subsampling

Subsamples from each mastodon were taken at their respective institutions and sent to the McMaster University Ancient DNA Centre. Subsamples were opened and processed in dedicated ancient DNA clean rooms thereafter.

### 2.2 Taxonomic identification of the Tualatin mastodon

The Tualatin mastodon is designated F-30282 in the University of Oregon Museum of Natural and Cultural History (UOMNCH), but much of the specimen is on display in the Tualatin Public Library and the Tualatin Historical Society. The skeleton is partially complete with most of the left side of the animal present. Portions of the crania were recovered but were either discarded or lost over time. However, a fragment of the left maxillary tooth row containing the M2 and M3 was saved. It was recovered in 1962 along the Tualatin River, a tributary of the Columbia River, Oregon. Previous research on the specimen reports an age of 13,114-13,706 years BP ^11^ (Table 1), which post-dates the Missoula flood events and human colonization of the Pacific Northwest ^12,13^. Following published methods (i.e. ^7,14^), the molars were measured using digital calipers. These measurements are compared to a large dataset of *Mammut* upper M3s, including *M. pacificus* (N=39; ^7^), interglacial *M. americanum* from Alaska and the Yukon (N=7; Clades A and Y) and late glacial *M. americanum* from the Great Lakes (N=46) (Supplementary Data – Table S1).

**Table 1.** Coverage statistics. Mapping and coverage statistics for the new mastodon specimens sequenced as part of this study. Coverage values represent percent coverage of the reference at 3X or higher, and the mean coverage across resolved positions. The depositional context in which the specimens were found, their associated (A) or direct (D) radiocarbon ages or electron spin resonance dates, as well as their common names are also provided.

| Specimen | Depositional Context | Specimen ID | # Mapped Reads | Mean Frag. Size (bp) | Coverage | Age (BP) | References |
| --- | --- | --- | --- | --- | --- | --- | --- |
| Tualatin | Swampland | UOMNCH F-30282 | 22,955 | 45.2 | 99.3% / 63.2x | 11,480 ± 35 (UCIAMS78127; <sup>14</sup>C - D) | 11 |
| Renison | Unknown | ROM41948 | 2,297 | 42.2 | 79.4% / 5.3x | - | 36 |
| Little Narrows | Gypsum Sinkhole | NSM015GF007 | 5,100 | 34.5 | 86.5% / 10.6x | 40,900 ± 2300 (UCIAMS157082; <sup>14</sup>C - D) | This study |
|  |  |  |  |  |  | >52,800 (UCIAMS157040; <sup>14</sup>C - A) | This study |
|  |  |  |  |  |  | >50,300 (UCIAMS157044; <sup>14</sup>C - A) | This study |
| Milford | Gypsum Sinkhole | NSM019GF009.002 | 260,585 | 52.2 | 100% / 827.2x | 74,900 ± 5000 (ESR - A) | 31 |
| Georges Bank | Submerged Terrestrial | NSM021GF005.001 | 62,834 | 49.1 | 100% / 187.7x | - | - |
| Middle River | Gypsum Sinkhole | NSM979GF153.001 | 8,558 | 41.8 | 98.2% / 21.7x | - | 37 |
| Windsor | Gypsum Sinkhole | NSM989GF089.002 | 21,458 | 35.3 | 99.1% / 46.1x | - | 37 |

A three-dimensional model of the Tualatin specimen was also created with Structure-from-Motion (SfM) photogrammetry. (Morphosource ID: 000683205). The specimen was photographed with a Nikon d5000 12.3 MP DX Digital SLR camera with a Nikkor 18-55mm f/3.5-5.6G VR II lens. The camera was mounted on a tripod positioned 40 cm from the specimen. The specimen was placed on a turntable and rotated 10 degrees for each photo while the camera was operated with a remote shutter release to reduce image blur. The specimen was photographed at three different angles (15, 30, and 45 degrees) to capture the entire surface. A total of 235 images were used to create a digital 3D model, generated using Agisoft Metashape Standard Version 1.7.1 software following established methods (e.g. ^15–17^).

### 2.3 New radiocarbon ages

A bone sample from specimen NSM015GF007 (Little Narrows) was sent for radiocarbon analysis to the Keck-CCAMS facility at the University of California, Irvine. The sample was decalcified using 0.5M HCl, rinsed with Milli-Q water, hydrolyzed overnight at 60°C with 0.01M HCl, and the high molecular weight fraction isolated.

Two wood samples associated with the collection site of NSM015GF007 were sent for radiocarbon analysis to the Keck-CCAMS facility at the University of California, Irvine. The samples were treated in 1N HCl at 70°C for 30 min to dissolve any contaminating carbonate from dust or soil, then washed with 1N NaOH at 70°C for 30 min to remove soil humics, repeating until clear, and finally washed with 1N HCl at 70°C for 30 min to remove atmospheric CO_2_ absorbed during the alkaline washes. Samples were washed with MQ water to remove chloride and dried on a heating block or in a vacuum oven.

### 2.4 DNA extraction and sequencing

Samples were initially processed as previously described ^4,9^ with a few modifications. Briefly, approximately 150-350mg of material was subsampled from each specimen and manually pulverized. Subsamples were prewashed with 300µl of 0.5M EDTA for 20min at 1000 RPM to remove dust, and the wash was discarded. Subsamples were then demineralized using 0.75-1ml of 0.5M EDTA pH 8.0 (room temperature; 2-5 days with shaking at 1000-2000 RPM) and digested using 0.75-1ml of a Proteinase K digestion buffer (0.01M Tris-CL (pH 9); 0.20% Sarcosyl; 0.25 mg/ml Proteinase K; 0.01M CaCl_2_; 45°C for ∼3 days with rotation) in two successive rounds. Demineralization and digestion supernatants were pooled for extraction. All processing steps were accompanied by an extraction blank which was treated identically to the subsamples but contained no material.

DNA was extracted as described in Dabney et al. ^18^ with the following modifications: Roche High pure viral nucleic acid columns were used instead of MinElute columns; 1ml of supernatant was mixed with 13ml binding buffer and successively passed through the column until all supernatant was processed; DNA was eluted twice with 25μl of EBT (Buffer EB + 0.05% Tween-20).

Extracted DNA was converted into non-UDG-treated libraries using a double-stranded library protocol. Double-stranded libraries used ∼20µl of input in 40µl reactions using previously described methods ^19,20^ with the following modifications: NE Buffer 2.1 replaced NE Buffer 2; MinElute clean-ups were done with two 750µl PE Buffer washes, and an additional 60s dry spin; libraries were heat deactivated at 80°C following adapter fill-in instead of column purified. 12.5µl of each library was used for indexing PCR with unique P5 and P7 index adapters. Indexing PCR was performed in 40µl reactions (1X KAPA SYBR Fast qPCR Master Mix; 750nM each P5/P7 indexing primer) for a max of 20 cycles (denaturing at 95°C for 30s; annealing at 60°C for 45s), although libraries were pulled earlier if they were observed to be undergoing exponential amplification. Indexing PCR reactions were then purified over MinElute columns as before, except eluted in 13µl of EBT.

To increase endogenous DNA, we also performed one round of in-solution enrichment using a previously published proboscidean mitochondrial genome bait set ^21^. Double-stranded libraries were processed as described previously ^9^ using 9.05µl of indexed library.

Enriched libraries were pooled to approximately equimolar concentrations and size selected for fragments between ∼150 and 500bp (3% Nusieve GTG Agarose Gel; 100V for 35 min). DNA was purified from gel plugs using the QIAquick Gel Extraction Kit with the following modifications: an additional 700µl PE Buffer wash; an additional dry spin at maximum speed; and elution in 20µl. Libraries were sequenced on an Illumina HiSeq 1500 using 2x90bp chemistry.

We subsequently extracted more material for the Renison and Tualatin mastodons as our initial sequencing did not generate complete mitochondrial genomes (>80% coverage of the reference genome at a minimum depth of 3x). Six additional extracts each were generated from the Renison molar and the Tualatin pelvis (Table S1) with ∼25-50mg of input. Material was demineralized as described in Dabney et al. ^18^ (for libraries ending in A or D in Table S1), or as above with a modified Proteinase K digestion buffer (0.02M Tris-CL (pH 9); 0.5% Sarcosyl; 0.25mg/ml Proteinase K; 0.005M CaCl2; 1% PVP; 50mM DTT; 2.5mM PTB) for 24 hours (for libraries ending in B, C, E, or F in Table S1). DNA was extracted as described above except eluted in 2x20µl in TET Buffer (0.01M Tris-HCL; 0.001M EDTA; 0.05% Tween-20).

All subsequent extracts were converted into single-stranded libraries using an established protocol ^22^. 12.5µl of each library was indexed as described before and subject to two rounds of in-solution enrichment as described above. Enriched libraries were pooled in equimolar concentrations and sequenced on a NextSeq 2000 using 2x50 bp chemistry.

### 2.4 Sequence processing and curation

Reads were mapped to the *M. americanum* mitochondrial reference genome (NC_035800) extended by ∼20 bp on either end as previously described ^9^. In brief, demultiplexed reads were trimmed and merged using the ancient DNA settings in leeHom ^23^. Reads were mapped to the padded reference with a network-aware version of BWA ^24^ (https://github.com/mpieva/network-aware-bwa) with established ancient DNA settings: maximum edit distance of 0.01 (-n 0.01), a maximum of two gap openings (-o 2), and seeding effectively disabled (-l 16500). Mapped reads which were merged or properly paired were extracted using the retrieveMapped_single_and_ProperlyPair program of libbam (https://github.com/grenaud/libbam), and collapsed based on unique 5′ and 3’ positions to remove PCR duplicates (https://bitbucket.org/ustenzel/biohazard/src/master/). Reads were then filtered to a minimum fragment size of 24bp and a minimum mapping quality of 30 using SAMtools ^25^. Specimens for which multiple libraries were sequenced were then combined using the merge function of SAMtools. To ensure authenticity we also examined the cytosine deamination signal in our final alignments using MapDamage2.0 ^26^.

Alignments were then imported into Geneious (v2021.2.2) and manually curated to remove sequencing artefacts and insertions not supported by a majority of the aligned reads. A consensus was called by strict majority, and any position with less than 3x coverage masked with Ns. We further masked a portion of the 16S region in the Renison mastodon alignment as it contained many short, stacked reads which have previously been shown to likely be of bacterial origin ^9^. The variable number tandem repeat region of each new consensus was also masked as in the NC_035800 reference, and the final consensus sequences were truncated from either end to account for the padded reference.

### 2.5 Phylogenetic analysis

Consensus sequences for the new mastodon specimens were aligned to all available complete mastodon mitochondrial genomes (n = 37), both with and without the partial Renison mitochondrial consensus, using Muscle v5.1 ^27^. Separately, we also aligned the full dataset using two mammoth mitochondrial genomes (NC_007596 and NC_015529) as outgroups in rooted phylogenies. Model selection and maximum likelihood phylogenies were conducted using IQ-TREE v1.6.12 ^28,29^, choosing the model which minimized the AICc estimate in each case. Maximum likelihood trees were generated with 1000 bootstraps and the chosen model: all mastodons (TPM3u+F+I+G4); all mastodons without Renison (TPM3u+F+I+G4); all mastodons with mammoth outgroups (TIM3+F+I+G4). We subsequently also constructed phylogenies with each of the datasets above using an HKY+G4+F model (used in subsequent analyses with BEAST) to verify the substitution model should not affect our observed topology.

BEAST analyses were conducted as per the Joint analysis described previously ^4,9^ in BEAST v1.15^30^. In short, the median calibrated age for each specimen with a finite radiocarbon date was supplied to calibrate the model. Specimens with unknown ages were fit with diffuse gamma distributions (shape = 1; scale = 200,000), and a uniform prior restricting their age using priors between 800 kya and 50 kya (for specimens which were/deemed to be non-finite) or 0 kya (for specimens which may potentially yield finite ages). NSM019GF009.002 (Milford mastodon) is a juvenile mastodons associated with the adult East Milford mastodon, N.S., and as such is also expected have a date of 74.9 ±5.0 ka ^31^, and a previously obtained radiocarbon estimate for the Tualatin mastodon ^11^ was calibrated in Calib v8.2 ^32^ to produce a median estimate of 13,360 yBP. Three newly obtained radiocarbon ages related to the Little Narrows mastodons (NSM015GF007) were consistent with this specimen being outside the limits of radiocarbon dating (Table 1; Tables S5-S6), and as such it was treated as other non-finite mastodons. The uniform bounds on Georges Bank mastodon (NSM021GF005.001) were set to between 0 and 800 kya due to this specimen’s position in Clade G which is comprised entirely of mastodons dated to <50 ky. We subsequently ran an additional model, wherein the ages of each mastodon, not generated as part of this study, were supplied as point estimates equal to their median calibrated radiocarbon age, or the median age that we previously estimated ^9^. Three independent chains were run for 500 million generations (sampling every 10,000) to assess convergence in each model. Chains were combined for the final analysis.

## 3. Results

### 3.1 DNA recovery and coverage statistics

We managed to recover six complete mitochondrial genomes from mastodon specimens in this study – five from Nova Scotia/the Georges Bank Region and one from Tualatin, Oregon (Fig. S1). We also managed to recover a partial mastodon mitochondrial genome from northern Ontario (Renison), for which we managed to reconstruct 79.4% of the *M. americanum* mitochondrial reference (NC_035800) at a mean depth of 5.3X (Table 1). All mastodons analyzed in this study had short mean fragment sizes (34.5bp - 52.2bp; Table 1) and characteristic deamination patterns consistent with authentic ancient DNA (Fig. S2). The extraction blank processed alongside the samples recovered <20 reads mapping to the conserved regions of the mitochondrial genome.

### 3.2 The Tualatin mastodon extends the range of Pacific mastodons

Recent studies highlighting morphological and genetic variability in late Pleistocene mammutids suggest that *M. americanum* has a more complex population history than previously appreciated ^4,7,9^. Currently, it is unclear if these population-level differences reflect species-level divergence. However, these differences reflect the reality of deeply divergent populations, part of a continent-wide metapopulation, which we refer to as the *M. americanum* complex. Consequently, we have compared molar L:W ratios obtained for the Tualatin mastodon and compared them against two regionally distinct groups of American mastodons.

The Tualatin mastodon M3 has a crown length (L) of 149.89 mm, a maximum crown width (W) of 78.61 mm, producing a L:W ratio of 1.91. This molar is small, but plots comfortably within the range for *M. pacificus* (Fig. 1). The L:W is within 1 standard deviation of the mean L:W of *M. pacificus*, although it is slightly outside the lower boundary of the 95% CI (1.94-2.02). It is well above the upper boundary of the *M. americanum* 95% CI (1.73-1.77). Based on its location within the morphospace defined by the *M. americanum* complex and *M. pacificus*, the Tualatin mastodon is confidently identified as *M. pacificus* ^7^. This result makes the Tualatin mastodon the first Pacific mastodon reported in Oregon. It also expands the known geographic range of the Pacific mastodon into the Pacific Northwest.

**Figure 1.**
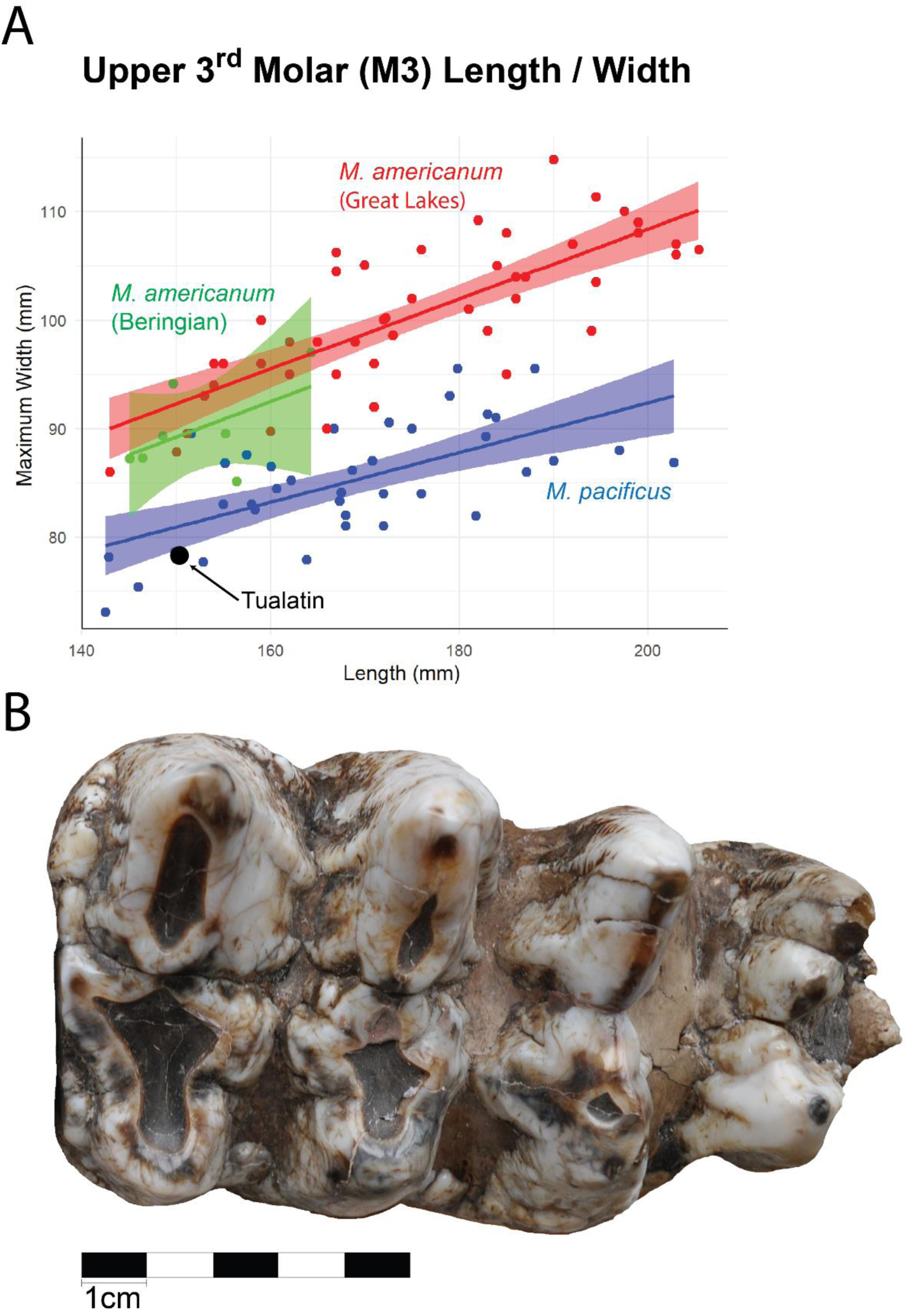
Tualatin mastodon. **A.** The relationship between molar length and maximum width in the upper third molar (M3) of the Tualatin mastodon in comparison to *M. pacificus* and *M. americanum* M3s from the Great Lakes region and Beringia. The 95% confidence interval around each regression line is indicated in the shaded colour. **B.** Upper occlusal view of the Tualatin mastodon’s M3.

Phylogenetically, the Tualatin mastodon from Oregon groups within Clade M, and most closely with a mastodon from central Alberta (RAM P97.7.1; Fig. 2). These two samples share a common ancestor ∼440 kya, although, given the uncertain age estimate on RAM P97.7.1, the 95% HPD interval on the node remains broad (230-690 kya). Importantly however, this age estimate falls near the palaeontological appearance of *M. pacificus* ^33^, and suggests that RAM P97.7.1, originally identified as an American mastodon, may likely be better reclassified as Pacific mastodon. This would extend the range of *M. pacificus* deep into western Canada, from its previous upper limit of Montana ^33^. Interestingly, the only other mastodon in Clade M is a late Pleistocene mastodon from central Mexico (DP1296). This specimen is deeply diverged and is occasionally paraphyletic with respect to other mastodons in Clade M (Fig. 2; see Figs S4-S9 and S11 for comparison). Resolving the exact relationship within this clade will require more dated mastodons. However, assignment of these other two members of this clade as *M. pacificus* suggests at least two possible scenarios; i) either that Mexican mastodon, DP1296, should also be considered a Pacific mastodon and the range of this species extends well outside of the contiguous United States (i.e. Alberta to Mexico); or ii) that DP1296 represents yet another as of yet unidentified “species-level” lineage of mastodons from the southern parts of the continent.

**Figure 2.**
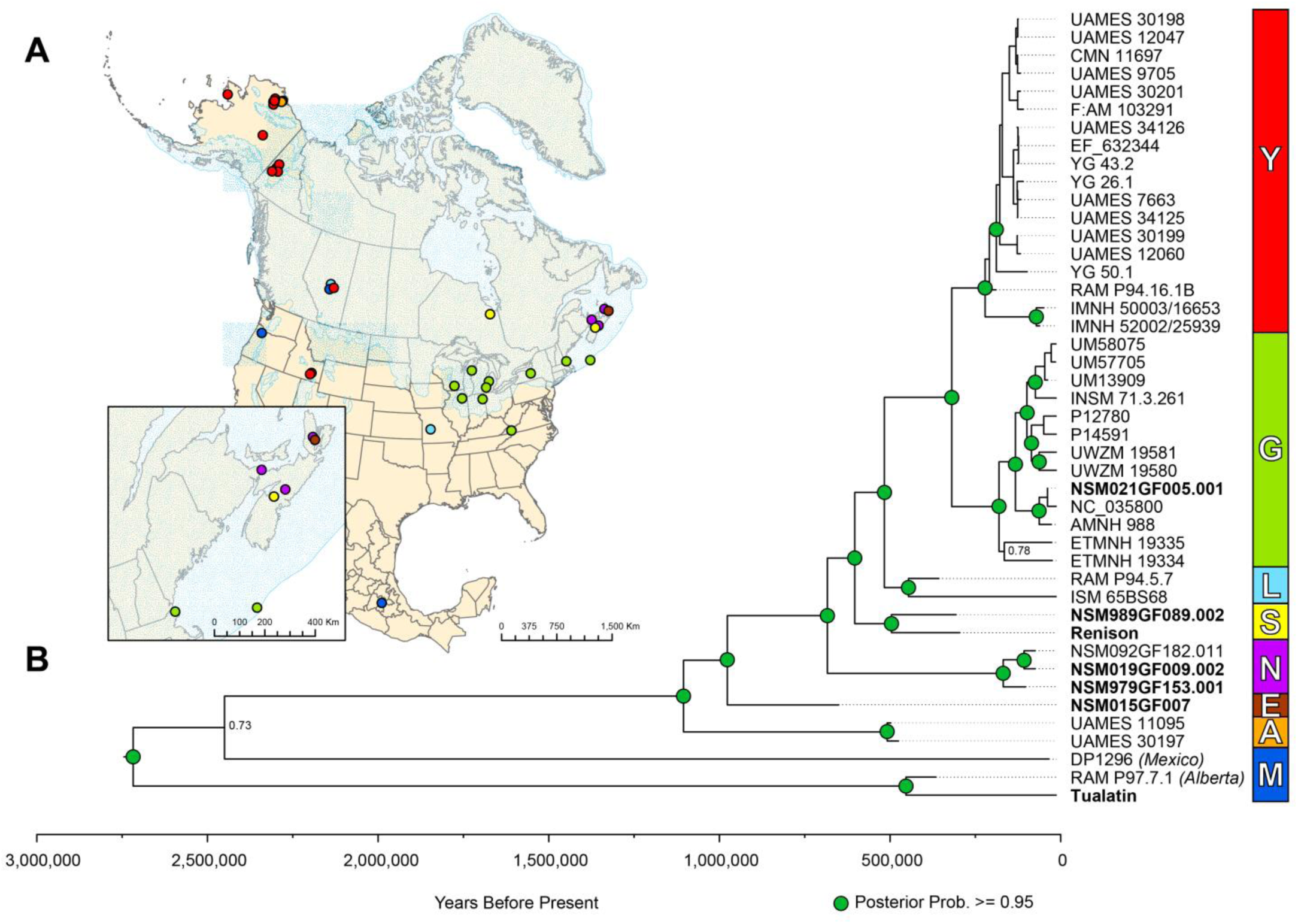
Mastodon phylogeography. **A.** Map of North America with all available complete mastodon mitochondrial genomes (and the partial genome from Renison) coloured as per their clade assignment. **B.** Maximum clade credibility tree generated when all undated/non-finite mastodons are fit with diffuse gamma distributions. Specimens generated as part of this study are shown in bold. Nodes with posterior probability values greater than or equal to 0.95 are indicated by green circles. Support for other key nodes is indicated in text next to the relevant node. The location of other mastodons in Clade M is shown in italics.

### 3.3 East coast mastodon mitochondrial genomes suggest at least three dispersal events

Our phylogenetic analysis of the five mitochondrial genomes from east coast mastodon samples and our partial mitochondrial genome from the Northern Ontario mastodon, has uncovered a surprising amount of diversity relative to the geographic proximity within which these animals were recovered. Interestingly only two of the new mastodon mitochondrial genomes fall within Clade N – the Milford (NSM019GF009.002) and Middle River (NSM979GF153.001) mastodons – which contained the only previously sequenced specimen from Nova Scotia (Fig. 2). The Georges Bank mastodon (NSM021GF005.001) clustered within Clade G, a clade comprised almost entirely of mastodons from the Great Lakes region. The remaining three clustered within two new clades – E and S. We do note that Clade E contains only a single specimen (Little Narrows; NSM015GF007) and as such is not a true clade, but it represents yet another deeply diverged group of mastodons.

We used Bayesian age estimation to date the samples and recovered signals of at least three temporally distinct groups in the eastern mastodon clades (Fig. 3). The Georges Bank mastodon produces a young age estimate (median: ∼38 kya; 95% HPD: 13-84 kya), consistent with its position in Clade G among other young mastodons that colonized the Great Lakes region and northeastern United States after the last glacial maxima. The estimated median age and posterior probability distribution of the Middle River mastodon (median: ∼103 kya; 95% HPD: 50-169 kya) is similar to other MIS 5 mastodons (∼71-130 kya), and close in age to the other two mastodons in Clade N, both of which are dated to 74,900 +/-5000 ky. The Windsor (NSM989GF089.002) and Renison (ROM41948) mastodons (median ages: ∼307 kya and ∼296 kya respectively) produce very diffuse distributions characteristic of very old mastodons from early time points in our phylogeny (Fig. 3). Although the bulk of the posterior probability is located at ages older than the 95% HPD range of Middle River, suggesting these mastodons are likely temporally distinct, there is some overlap between the lower range of these two mastodons and the Middle River mastodon (Windsor: ∼4.6 ky of overlap; Renison: ∼25 ky of overlap). Little Narrows (NSM015GF007), the sole representative of Clade E, also produces a diffuse age estimate, however, this distribution was highly concentrated on very old ages, abutting the upper bound we fixed on our analysis (median – ∼650 kya; 95% HPD: 464-800 kya). Notably, the posterior probability recovered for Little Narrows suggests it may be the oldest mastodon in our analysis (and the oldest mastodon sequenced to date) and has a non-overlapping 95% HPD with other east coast mastodons, suggesting it is temporally unique. Interestingly, this also highlights the stellar preservation potential of gypsum sinkholes, as this mastodon comes from a non-permafrost environment and is likely several hundreds of thousands of years old.

**Figure 3.**
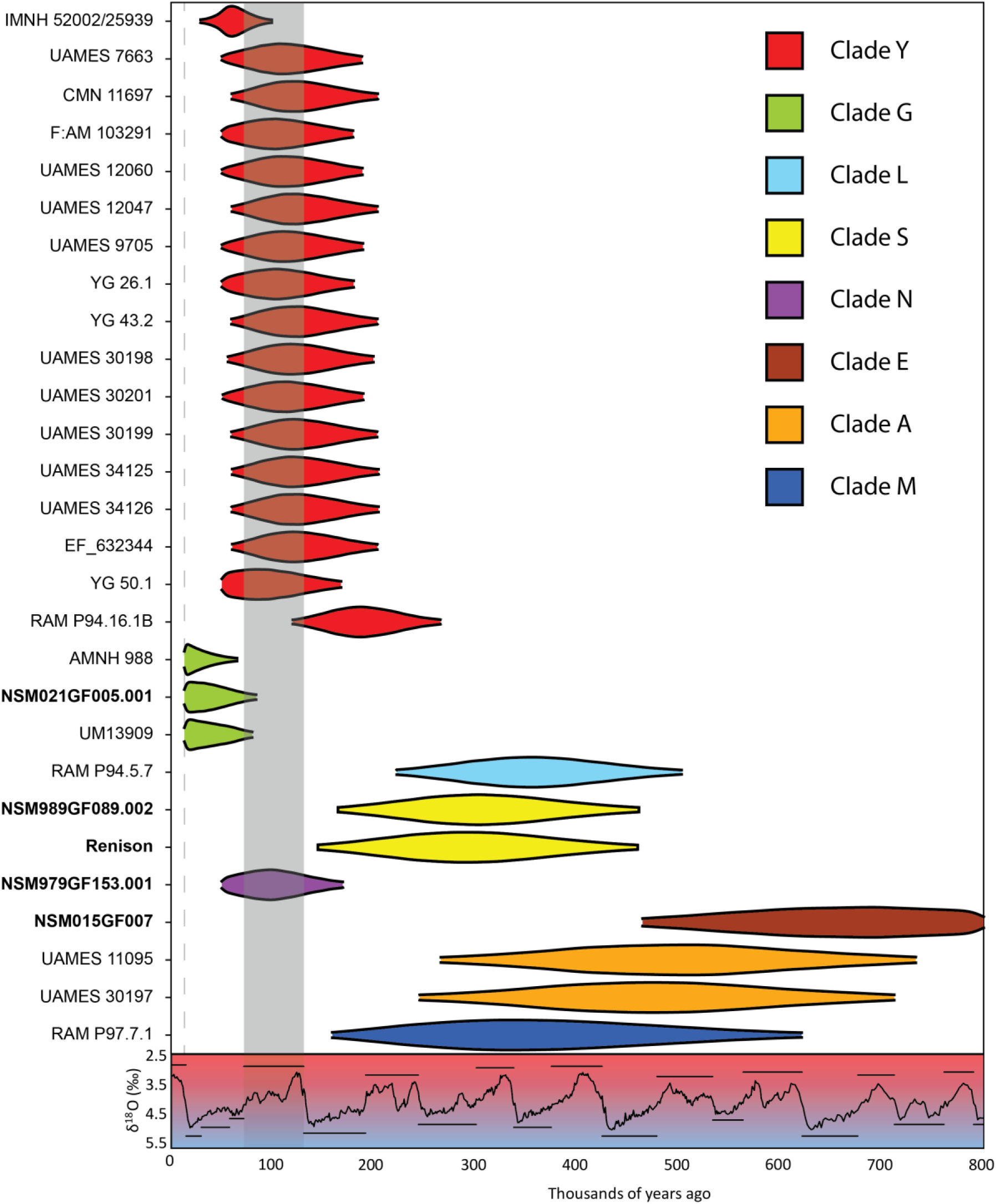
Mastodon age estimates. Marginal posterior densities of specimen ages when all unknown/nonfinite mastodons are fit with diffuse gamma priors. Each violin represents the 95% HPD interval estimated for each specimen. Specimens are coloured based on their clade assignments. Specimens generated as part of this study are bolded. The δ^18^O record for the last 800 thousand years ^38^ is overlaid below the plot, and the approximate extent of the MIS 5 interglacial highlighted in grey. The dashed line represents the lower bound of the analysis – 13,087 kya, the age of the youngest specimen (UWZM 19580).

We also examined the extent to which the amount of missing data in our analysis influenced our age estimates. To do this, we reran the previous BEAST analysis but supplied the median estimated age of specimens generated in previous studies as a point prior. We observed similar estimated median ages for each of our newly generated mastodon genomes (Table S4) and the same general patterns in the overlap of 95% HPD intervals, except the mastodons in Clade S no longer overlaps the 95% HPD of NSM979GF153.001 (Middle River; Clade N) (Table S4; Fig. S13). In general, we almost always recover smaller 95% HPD intervals, although this effect was most pronounced for our oldest specimen (Little Narrows; NSM015GF007), which saw an ∼30% reduction in the value range.

## 4. Discussion

This work expands on our understanding of *Mammut* taxonomy and biogeography throughout the middle and late Pleistocene in two substantive ways: i) we have generated a mitochondrial genome and its contextualization from an individual exhibiting morphology typical of *M. pacificus* and placed it within the broader diversity of North American mastodons; and ii) have produced a detailed investigation into the range dynamics of mastodons in the east coast of the continent. We have produced a model summarizing the phylogeography of Pleistocene *Mammut* in Figure 4.

**Figure 4.**
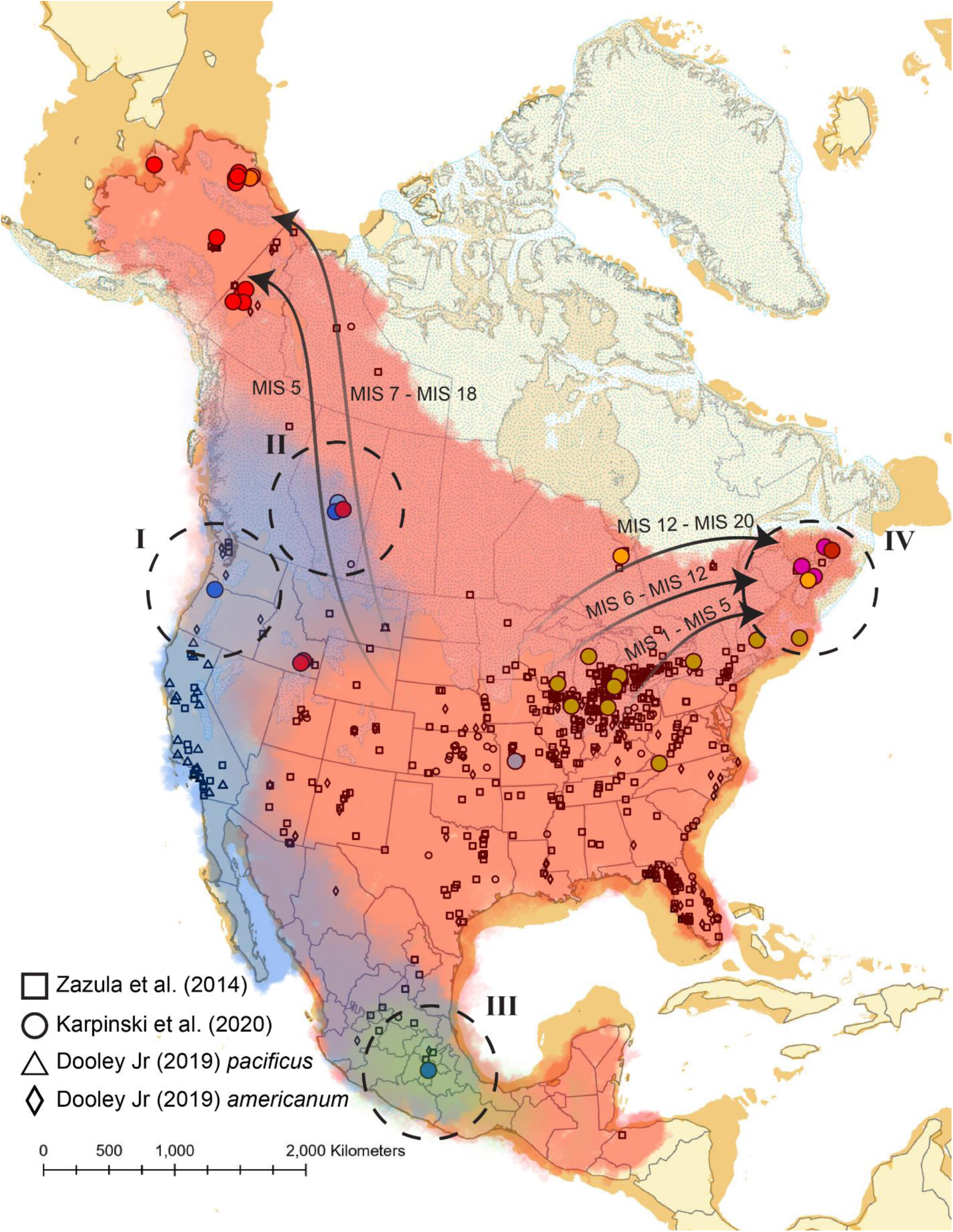
Pleistocene *Mammut* biogeography. Geographic model of our understanding of mastodon distributions and relationships throughout the mid-to-late Pleistocene. The approximate inferred distributions of the American mastodon morphological complex are shaded in red, and blue for the Pacific mastodon. The location of mastodon remains from previous large studies ^4,7,39^ are also provided (Supplementary Data – Table S2). The single *M. pacificus* specimen from ref ^33^ was combined with other Pacific mastodons from Dooley Jr. ^7^. Arrows indicate minimum confirmed expansions in response to glacial/interglacial cycles, with their timing ranges indicated in text. Arrows fade as they enter the contiguous United States due to the lack of a known source population. Key contributions from this study are indicated by dashed circles and roman numerals. **I.** The first mitochondrial data from a clearly identified Pacific mastodon, from Tualatin, Oregon. **II.** The reclassification of an Alberta mastodon (RAM P97.7.1) as *M. pacificus*, extending the range of Pacific mastodons into western Canada, and into regions where it might have been concurrent with mastodons showing characteristics of the *M. americanum* morphological complex. **III.** The possible extension of *M. pacificus* into Mexico, or the identification of another *Mammut* “species” (shown in green). **IV.** Increased genetic resolution in eastern North America, and the identification of at least 3 expansions into the eastern seaboard.

Previous work has presented conflicts between the palaeontological and genetic classification of American and Pacific mastodons ^9^ and suggested that the two species are not distinct at the mitochondrial level or that they exhibit similar patterns as seen between woolly and Columbian mammoths or extant bison (*Bison bison*), steppe bison (*B. priscus*), and giant long-horned bison (*B. latifrons*) ^34,35^. However, the study was complicated by the fragmentary nature of the samples, from which we recovered DNA, precluding morphological assessment as *M*. *pacificus* ^9^. In the current study, the completeness of the Tualatin mastodon and its unambiguous morphology have allowed us to clearly establish it as *Mammut pacificus*. Combined with its mitochondrial genome, this has allowed us to resolve a key question vis-a-vis mastodon taxonomy (Fig. 4 - I).

The phylogenetic position of the Tualatin mastodon (and possibly other pacific mastodons) within Clade M, may partially explain the high diversity of this clade relative to other clades, and extends the geographic range of pacific mastodons into Alberta, Canada (Fig. 4 - II). Additionally, morphologically and genetically well characterized American mastodons are known from the same geographic area as RAM P97.7.1 and with similar temporal estimates. This suggests these species likely interacted in Alberta and opens the possibility that they interbred, likely during northward expansions, as has been reported in black bears ^2^.

This study however raises additional questions surrounding the relationship of DP1296 (Mexico) to northern Pacific mastodons. Are we capturing a highly diverged lineage within *M. pacificus* or *M. americanum*, or rather have we identified a third, potentially cryptic species within North America (Fig. 4 - III)? Previous studies have shown that the diversity contained in Clade M is greater than within or between any of the other mastodon clades (see ref ^4^; Fig. S24). This is clearly represented in our phylogenies, where the branches separating the two lineages within Clade M are much longer than those within any of the other mastodon clades. While a reinterrogation of *Mammut* taxonomy and morphological variation is clearly needed ^9^, our results suggest that if *M. pacificus* continues to be considered a separate *Mammut* species, DP1296, and other associated Mexican mastodons, would be best considered a yet undescribed third species in Pleistocene North America.

Our east coast mastodon mitochondrial genomes reveal a greater amount of diversity not captured in previous studies, enabling us to examine the phylogeography of this region in much more detail. We observe at least three, but likely four, pulses into the region spanning ∼800 ky (Fig. 4 - IV). While clades E and N currently only include mastodons from Nova Scotia, given the geographic spread observed in clades S (Nova Scotia to northern Ontario) and G (the American Midwest to Northeast), it’s likely that these animals were part of much larger and highly diverse populations that expanded tracking shifting climates. However, except for the Middle River (NSM979GF153.001; Clade N) and Georges Bank mastodons (NSM021GF005.001; Clade G), we are unable to link these clades to specific glacial/interglacial periods during which these expansions might have occurred, as the posterior probability estimates are too diffuse and span multiple Marine Isotope Stages. Despite this lack of deep-time precision, this study contains the second such evidence of northward expansion and extirpation patterns in North American Pleistocene browsers and illustrates that these dispersal patterns were likely a regular occurrence on both sides of the continent in all transiently glaciated regions, not just Beringia.

This work highlights the importance of remains from outside the core, high-density ranges of mastodon species. Additionally, we show that even a few specimens from these areas can be immensely informative to key phylogeographic questions if chosen in a very targeted manner. This will likely be especially true for mastodons from more southern locales where biomolecular preservation is much worse, but where glacial refugia likely persisted (e.g. Florida) or which may shed light on the relationship between northern *M. pacificus* samples and their potential southern relatives in Mexico. There is also a need to obtain genetic data from mastodons with associated chronological information, outside of their late Pleistocene core range from both species to help calibrate phylogenetic analyses and reduce the uncertainty of our current temporal estimates and allow for more accurate linking between dispersals and major climatic events.

## 5. Author contributions

EK, HP, and CW conceived the study with feedback from AB and TF. AB preformed morphological analysis of the Tualatin specimen. TF sampled and collated information on the Nova Scotia material. EK and SB conducted all ancient DNA experiments. EK completed all bioinformatic analyses. EK wrote the first draft of the manuscript. All authors helped in the revision of subsequent drafts.

## 6. Data availability

Bulk sequencing data was deposited to NCBI’s SRA under bioproject PRJNA1190488. New mastodon mitochondrial genomes generated during this study were deposited to GenBank (PQ672301-PQ672307).

## 8. Acknowledgments

We would like to thank the Tualatin Ice Age Foundation, Tualatin Heritage Cernt, Tualatin Public Library, Yvonne Addington, Cindy Leigh, Jerianne Thompson, Mike Full, and the University of Oregon Museum of Natural and Cultural History for access and permission to sample the Tualatin mastodon, as well as the Nova Scotia Museum for access to the east coast mastodons, and to the Royal Ontario Museum for permission to sample the mastodon from Renison, ON. We would also like to acknowledge that the Little Narrows specimen (NSM015GF007) was found by quarry worker Sandy MacLeod and collected for the Nova Scotia Museum by Dr. John Calder and Katherine Ogden under permit P2014NS03. Likewise, The Windsor mastodon NSM989GF089.00 was found by quarry worker Alan Wilcox, and collected by Robert Grantham, Curator of Geology of the Nova Scotia Museum. We would also like to thank Brian Golding as well as HMS Research Computing for access to computational resources used in this work. We would also like to thank Brian Golding, Ben Evans, and all members of the Poinar, Golding, and Evans lab groups for support and feedback during this project. This work was supported by an NSERC Discovery Grant (grant No. 4184-15), and CRC to H.P.

