## Supplementary Methods/Figures for "Repeated climate-driven dispersal and speciation in peripheral populations of Pleistocene mastodons"

**Supplementary Table 1.** Specimen extract and library information for specimens analyzed in this study.

| Specimen | Sample Type | Mg Subsampled | Library | Library Methodology |
|---|---|---|---|---|
| NSM979GF153.001 | Femur | 196.6 | L1151 | Double-stranded |
| | | 160.5 | L1152 | Double-stranded |
| NSM989GF089.002 | Tusk | 231.5 | L1153 | Double-stranded |
| | | 343 | L1154 | Double-stranded |
| NSM019GF009.002 | Unknown Bone | 238 | L1155 | Double-stranded |
| | | 203 | L1156 | Double-stranded |
| NSM015GF007 | Bone Fragment | 217 | L1157 | Double-stranded |
| | | 314.5 | L1158 | Double-stranded |
| NSM021GF005.001 | Tooth | 201 | L1159 | Double-stranded |
| | | 211 | L1160 | Double-stranded |
| Renison | Molar | 234 | L1161 | Double-stranded |
| | | 286 | L1162 | Double-stranded |
| | | 52.9 | L7326A | Single-stranded (SCR) |
| | | 51.9 | L7326B | Single-stranded (SCR) |
| | | 25.3 | L7326C | Single-stranded (SCR) |
| | | 51 | L7326D | Single-stranded (SCR) |
| | | 52.7 | L7326E | Single-stranded (SCR) |
| | | 25.2 | L7326F | Single-stranded (SCR) |
| Tualatin | Molar | 183 | L1163 | Double-stranded |
| | | 185.5 | L1164 | Double-stranded |
| | Pelvis | 266 | L1165 | Double-stranded |
| | | 287 | L1166 | Double-stranded |
| | | 52.9 | L7320A | Single-stranded (SCR) |
| | | 50.7 | L7320B | Single-stranded (SCR) |
| | | 25.5 | L7320C | Single-stranded (SCR) |
| | | 52.5 | L7320D | Single-stranded (SCR) |
| | | 51.8 | L7320E | Single-stranded (SCR) |
| | | 25.2 | L7320F | Single-stranded (SCR) |

**Supplementary Figure 1.** Map showing the location of mastodons analyzed in this study. Newly sequenced mastodons are shown as circles, colored in red (American mastodon morphology) or blue (Pacific mastodon morphology). Pink crosses are used to indicate previously sequenced mastodons, while white diamonds indicate specimen locations from which DNA was not successfully extracted in a previous study. Mastodon locations have been jittered on the continental map to aid visualize.

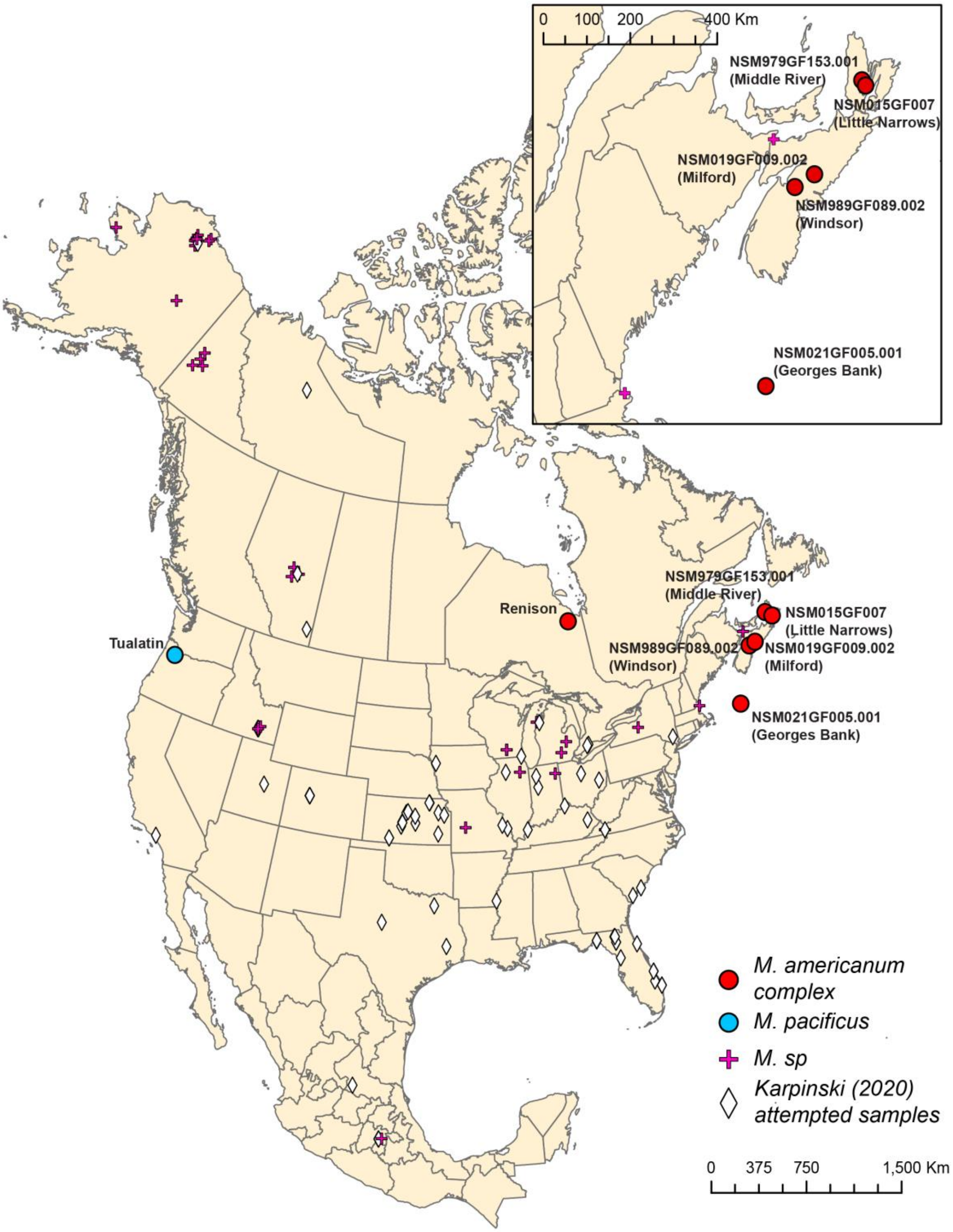

**Supplementary Figure 2.** Map Damage Plots for the final combined alignments for each specimen.

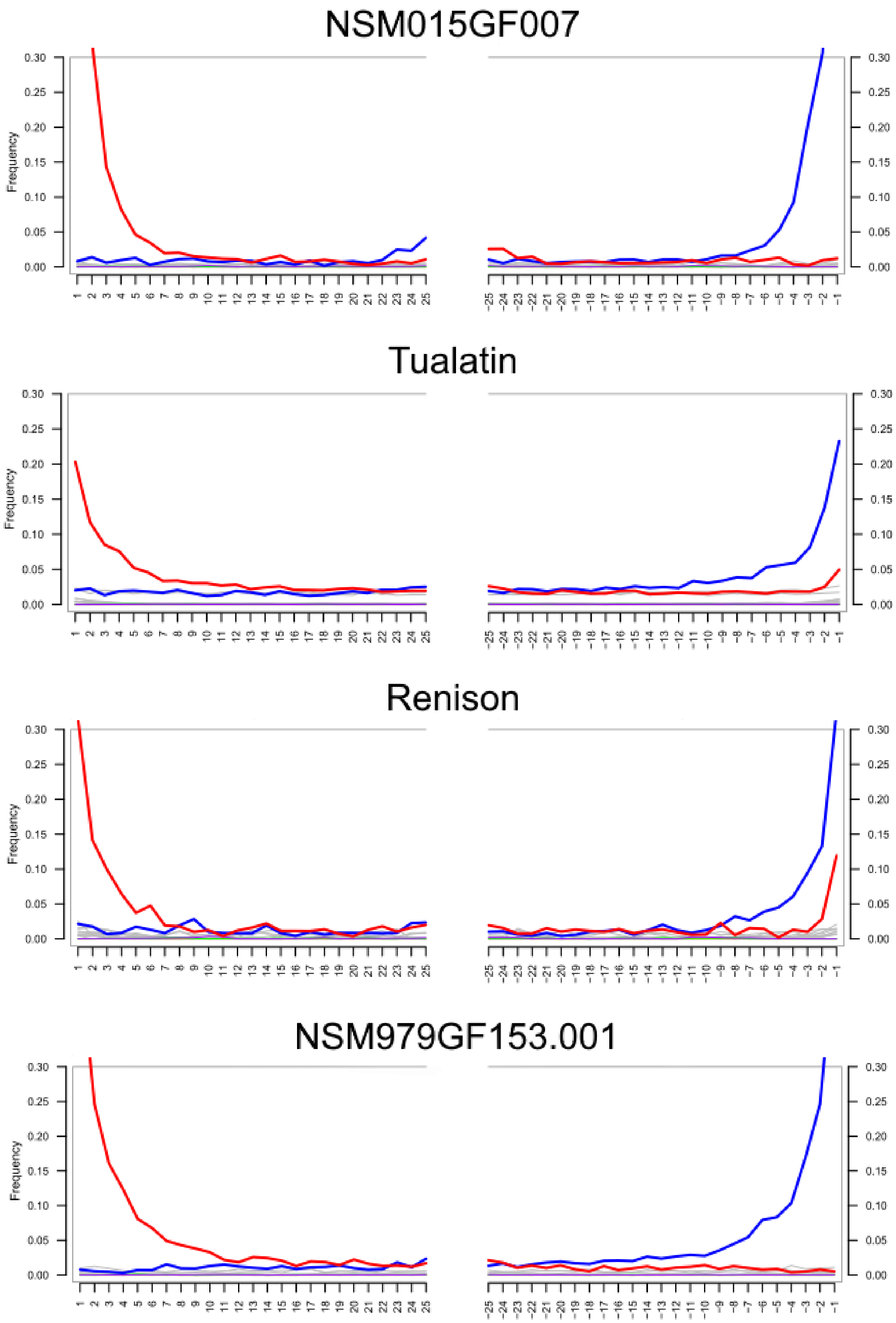

NSM989GF089.002

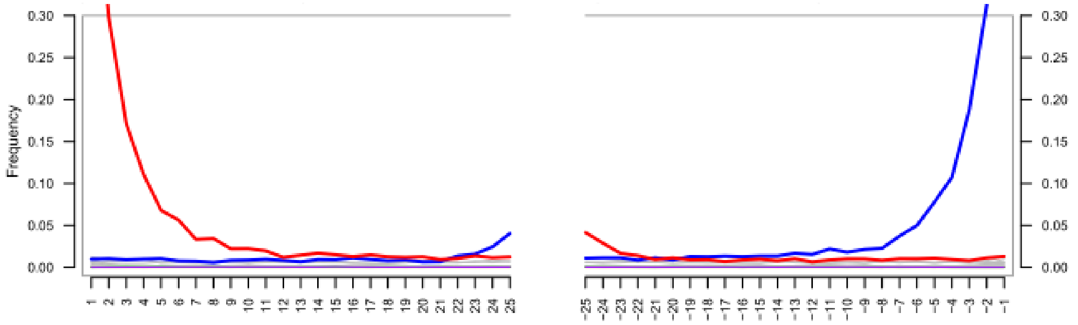


NSM019GF009.002

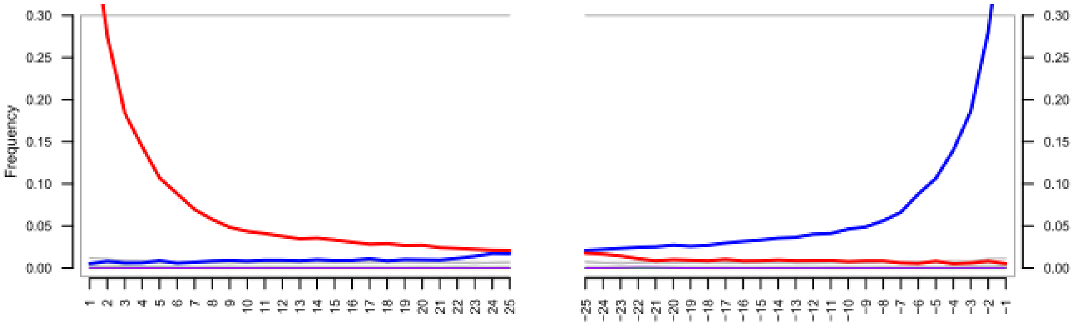


NSM021GF005.001

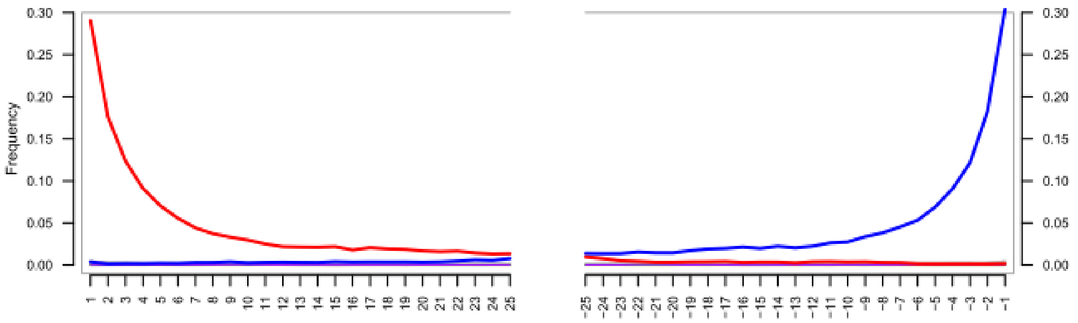

**Supplementary Figure 3.** Fragment length distributions of mapped reads for each specimen post sequence curation.

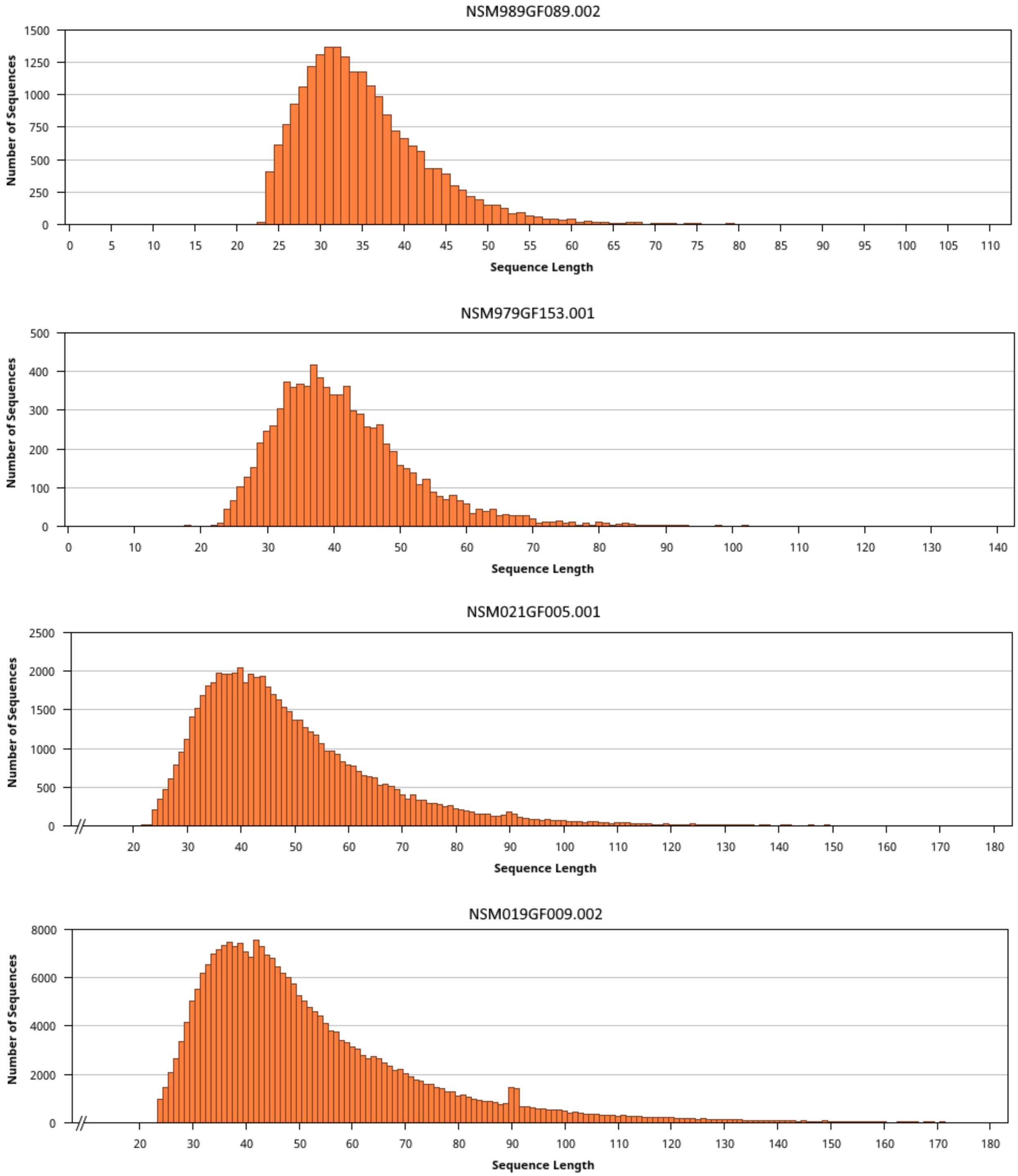

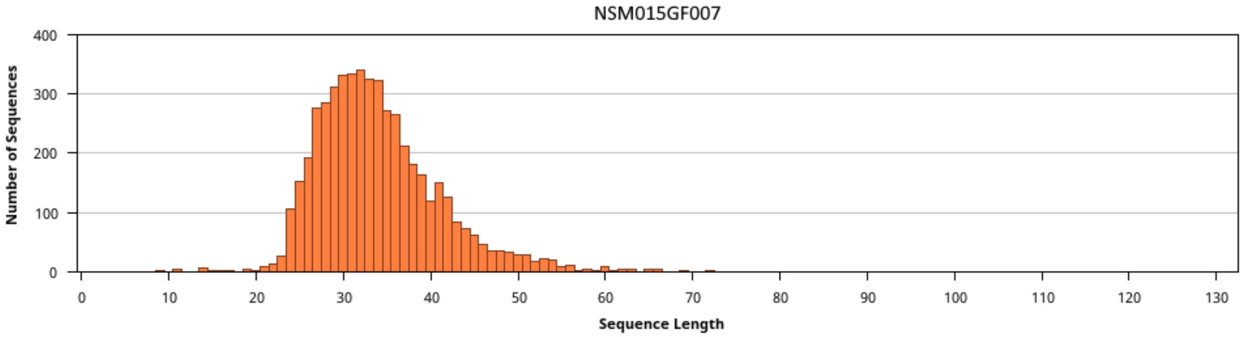
NSM015GF007
Number of Sequences
0
100
200
300
400
0
10
20
30
40
50
60
70
80
90
100
110
120
130
Sequence Length

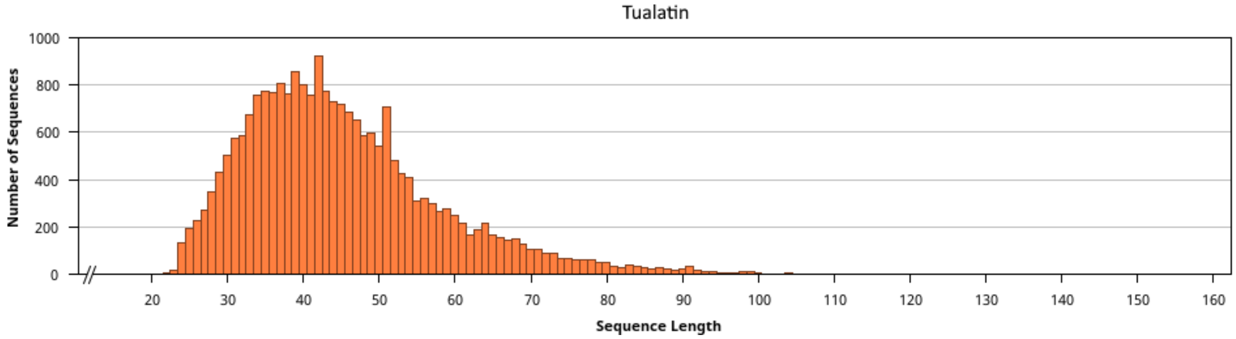
Tualatin
Number of Sequences
0
200
400
600
800
1000
20
30
40
50
60
70
80
90
100
110
120
130
140
150
160
Sequence Length

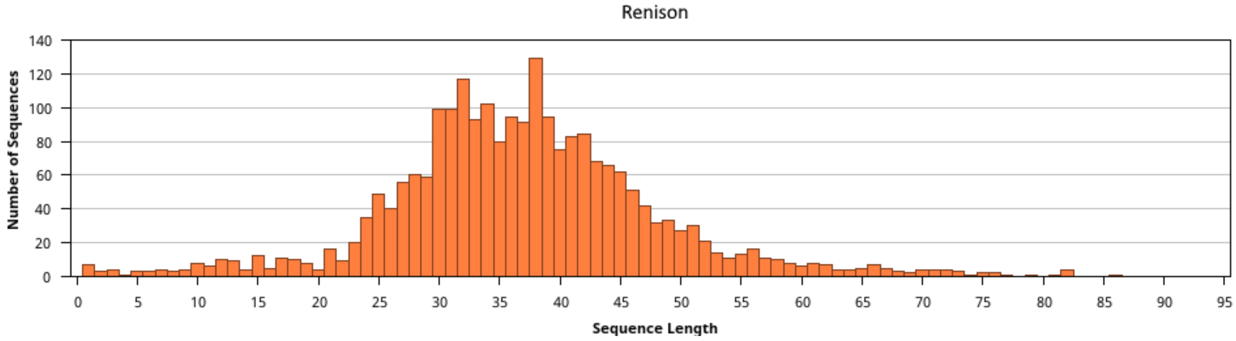
Renison
Number of Sequences
0
20
40
60
80
100
120
140
0
5
10
15
20
25
30
35
40
45
50
55
60
65
70
75
80
85
90
95
Sequence Length

**Supplementary Figure 4.** Midpoint-rooted maximum likelihood tree of all complete mastodon mitochondrial genomes plus the partial Renison sequence using the best fitting model (TPM3u+F+I+G4). Bootstrap values, shown at nodes, were assessed with 1000 bootstrap replicates.

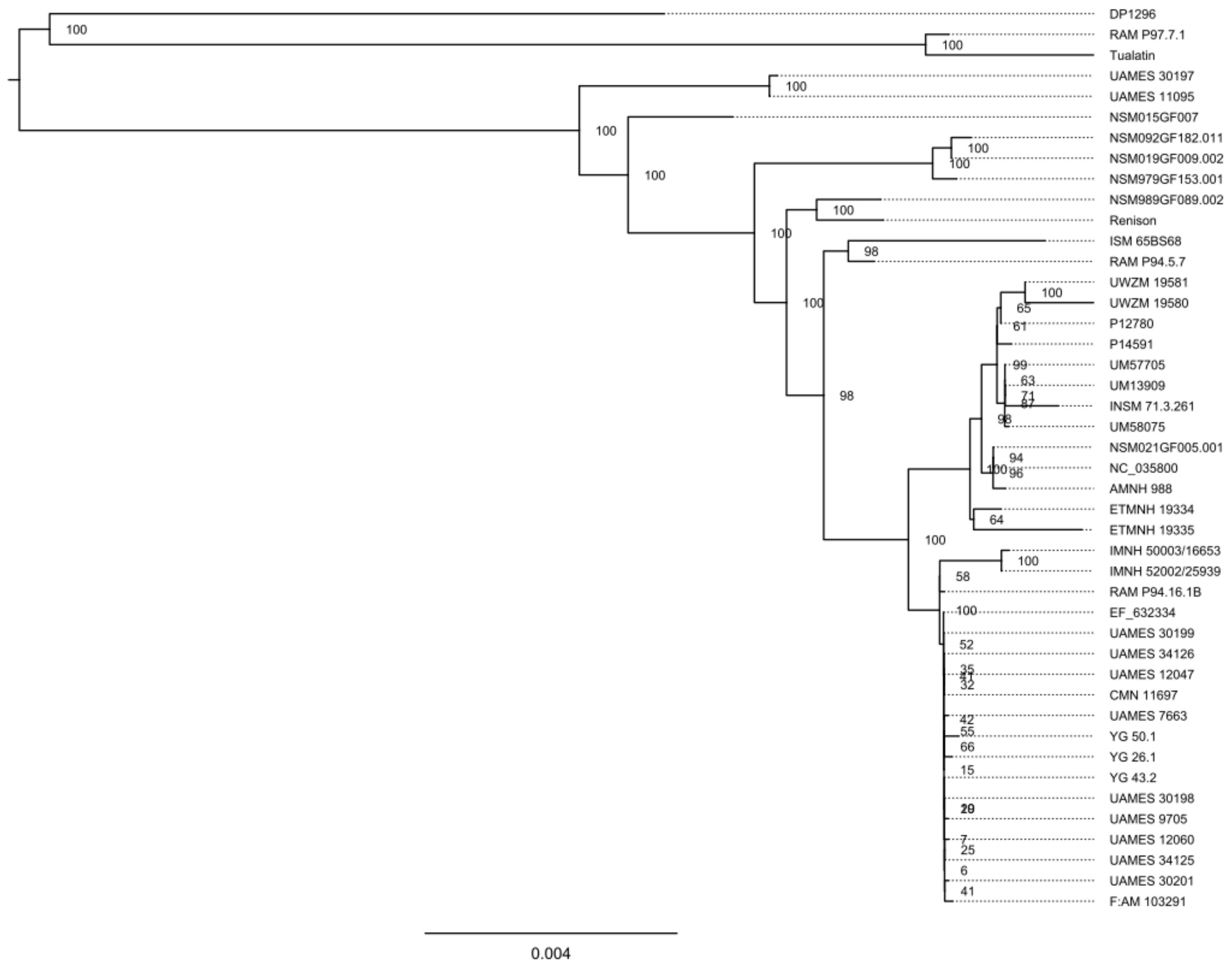

**Supplementary Figure 5.** Midpoint-rooted maximum likelihood tree of all complete mastodon mitochondrial genomes without the partial Renison sequence using the best fitting model (TPM3u+F+I+G4). Bootstrap values, shown at nodes, were assessed with 1000 bootstrap replicates.

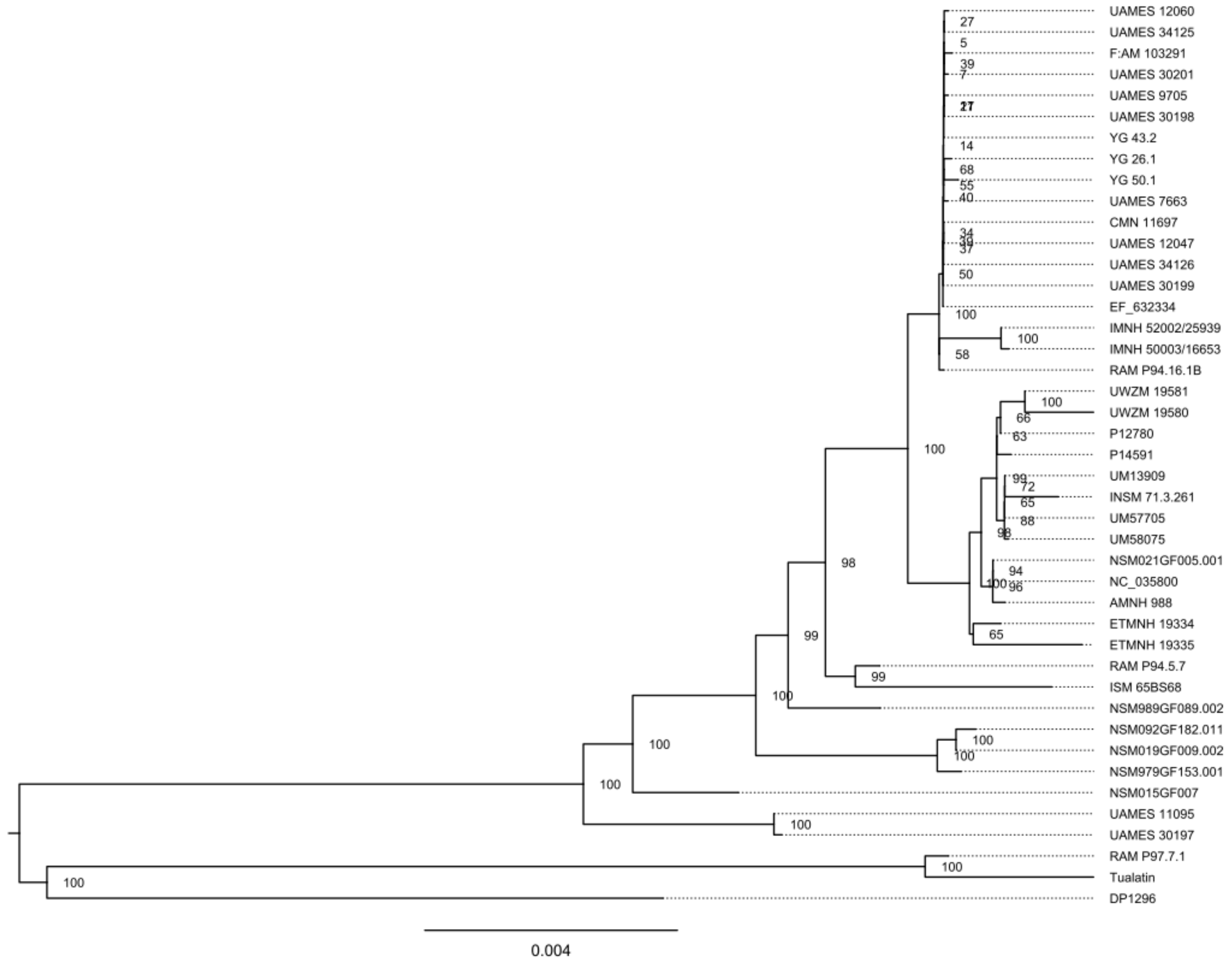

**Supplementary Figure 6.** Maximum likelihood tree of all complete mastodon mitochondrial genomes plus the partial Renison sequence and two woolly mammoth sequences using the best fitting model (TIM3+F+I+G4). Phylogeny was rooted using the woolly mammoth branch as an outgroup. Bootstrap values, shown at nodes, were assessed with 1000 bootstrap replicates.

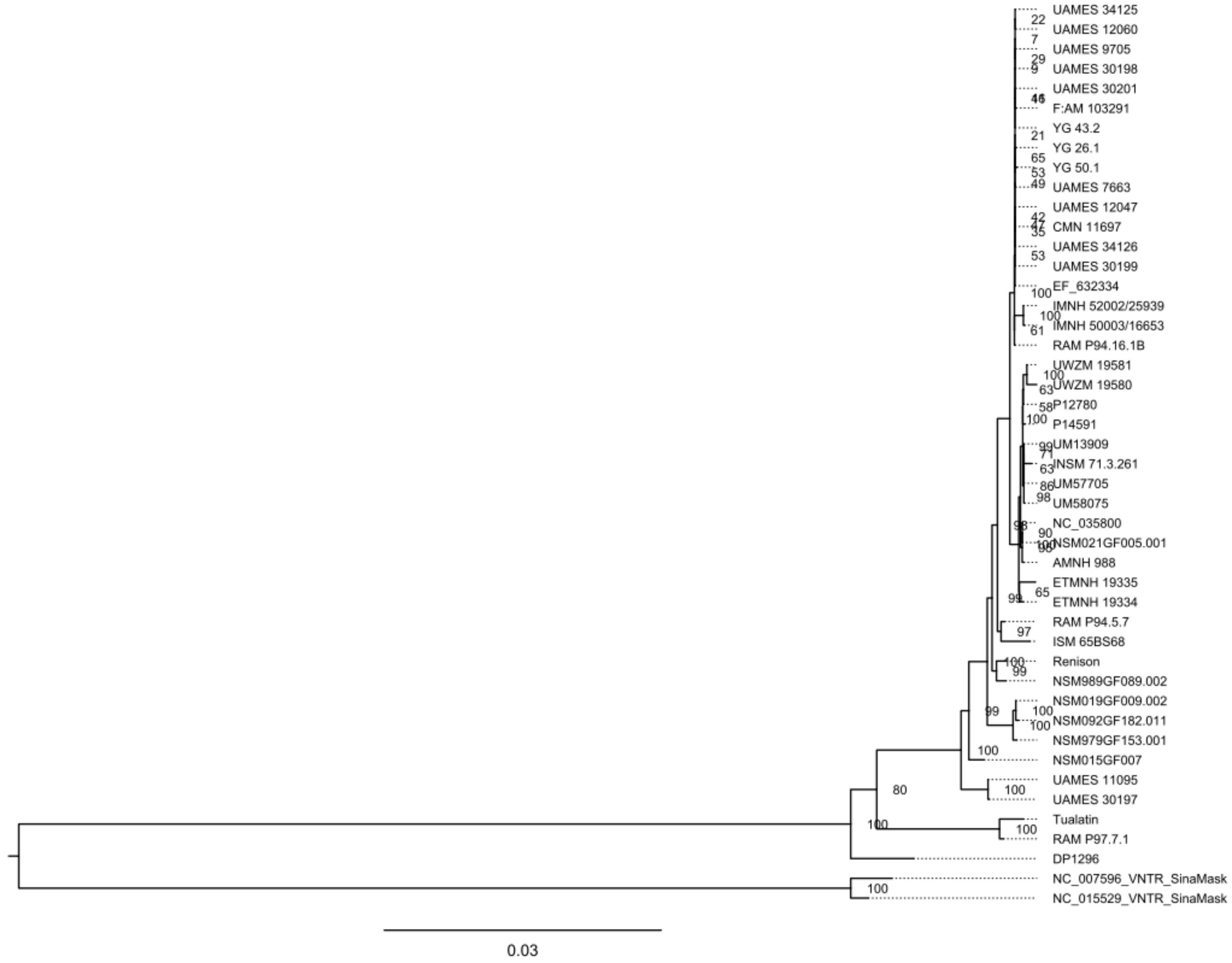

**Supplementary Figure 7.** Midpoint-rooted maximum likelihood tree of all complete mastodon mitochondrial genomes plus the partial Renison sequence using an HKY+G4+F model (used during Bayesian models). Bootstrap values, shown at nodes, were assessed with 1000 bootstrap replicates.

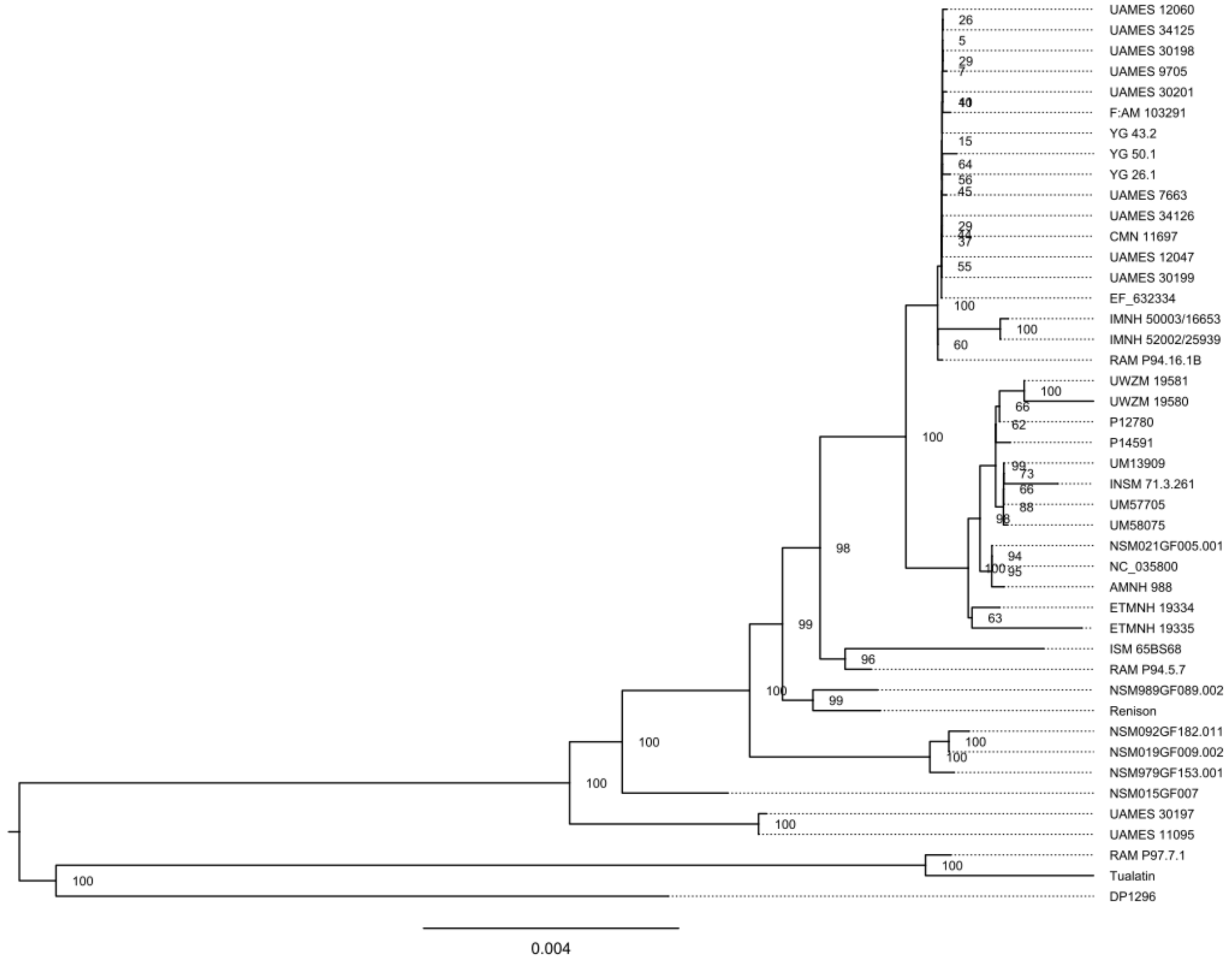

**Supplementary Figure 8.** Midpoint-rooted maximum likelihood tree of all complete mastodon mitochondrial genomes without the partial Renison sequence using an HKY+G4+F model (used during Bayesian models). Bootstrap values, shown at nodes, were assessed with 1000 bootstrap replicates.

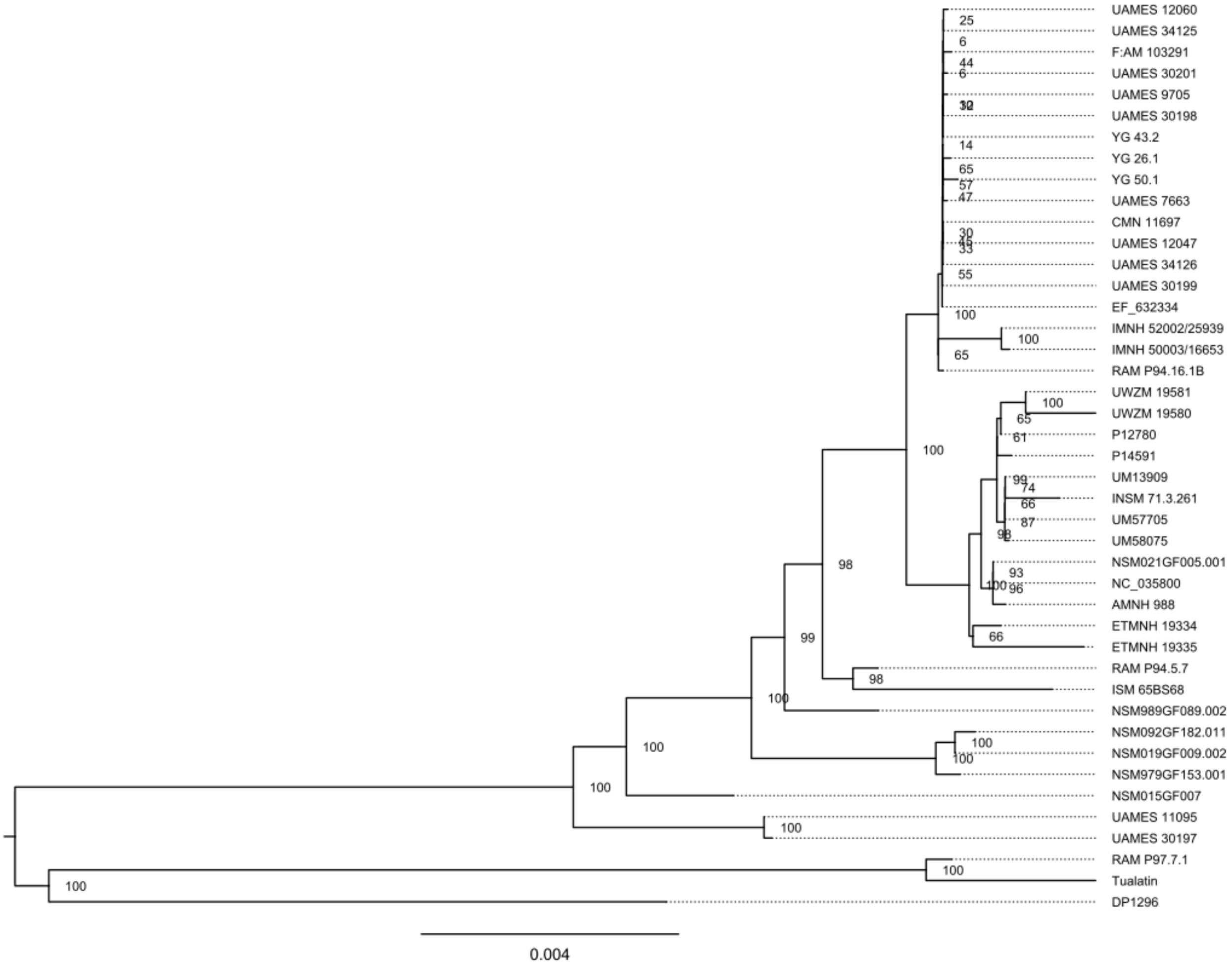

**Supplementary Figure 9.** Maximum likelihood tree of all complete mastodon mitochondrial genomes plus the partial Renison sequence and two woolly mammoth sequences using an HKY+G4+F model (used during Bayesian models). Phylogeny was rooted using the woolly mammoth branch as an outgroup. Bootstrap values, shown at nodes, were assessed with 1000 bootstrap replicates.

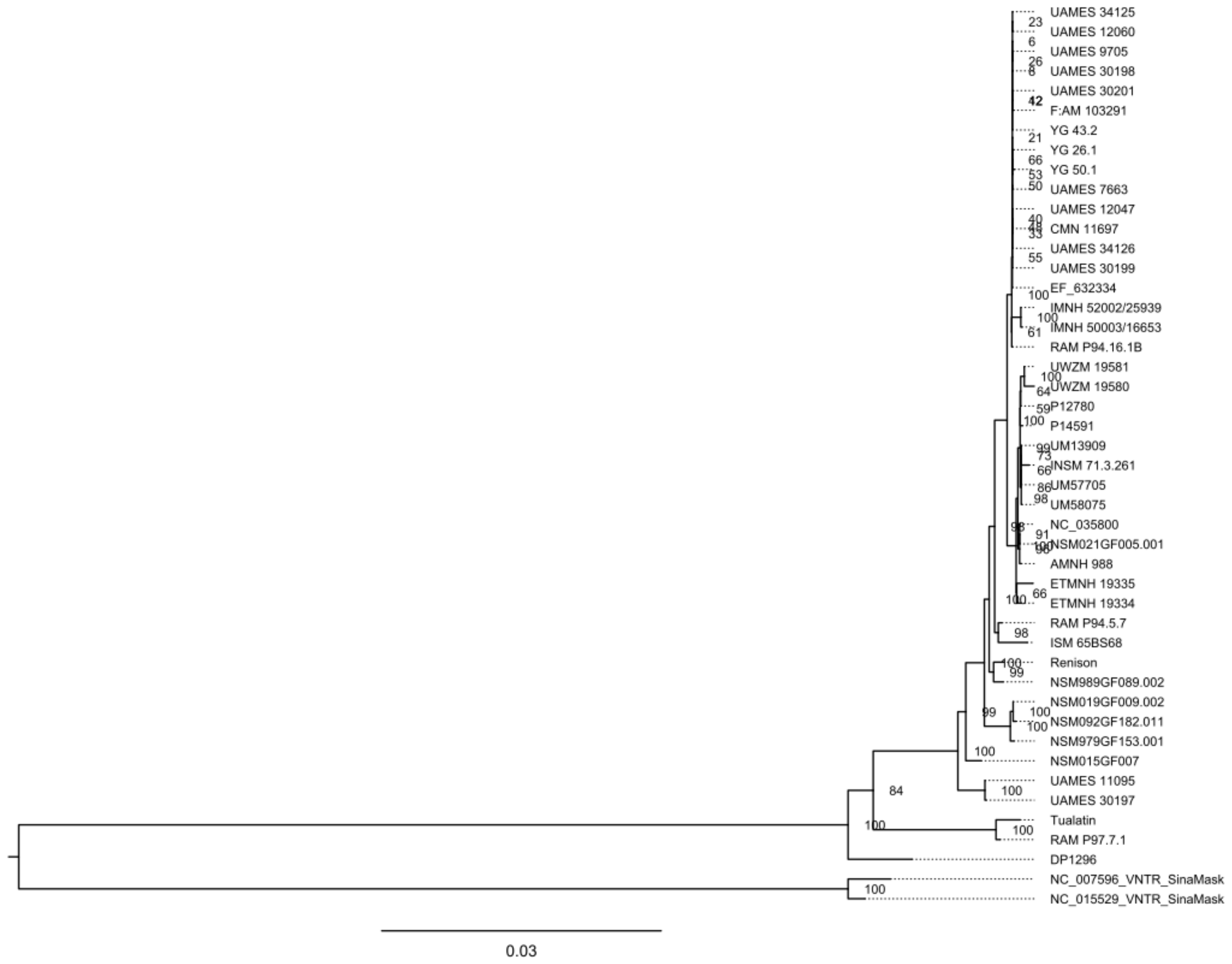

**Supplementary Table 2.** Parameters set on ages during BEAST analyses when sample was treated as an unknown or when the median estimate was used (values from Supplementary Table 5 of Karpinski et al. (2023)).

| Specimen | Unknown Age Analysis | Fixed Median Age Analysis |
|---|---|---|
| UAMES 12047 | Gamma:<br>- Shape: 1<br>- Scale: 200,000<br>- Offset: 0<br>Uniform:<br>- Lower: 36,913<br>- Upper: 786,913<br>Initial: 100,000 | Point estimate: 134,087 |
| NSM092GF182.011 | Point estimate: 74,900 | Point estimate: 13,488 |
| UAMES 30201 | Gamma:<br>- Shape: 1<br>- Scale: 200,000<br>- Offset: 0<br>Uniform:<br>- Lower: 36913,000<br>- Upper: 786,913<br>Initial: 100,000 | Point estimate: 125,087 |
| AMNH 988 | Gamma:<br>- Shape: 1<br>- Scale: 200,000<br>- Offset: 0<br>Uniform:<br>- Lower: 0<br>- Upper: 786,913<br>Initial: 20,000 | Point estimate: 27,714 |
| CMN 11697 | Gamma:<br>- Shape: 1<br>- Scale: 200,000<br>- Offset: 0<br>Uniform:<br>- Lower: 36913,000<br>- Upper: 786,913<br>Initial: 100,000 | Point estimate: 134,087 |
| P 12780 | Point estimate: 13,488 | Point estimate: 13,488 |
| UAMES 34126 | Gamma:<br>- Shape: 1<br>- Scale: 200,000<br>- Offset: 0<br>Uniform:<br>- Lower: 36913,000<br>- Upper: 786,913<br>Initial: 100,000 | Point estimate: 135,087 |

| | | |
|---|---|---|
| UM13909 | Gamma:<br>- Shape: 1<br>- Scale: 200,000<br>- Offset: 0<br>Uniform:<br>- Lower: 0<br>- Upper: 786,913<br>Initial: 20,000 | Point estimate: 41,977 |
| UAMES 11095 | Gamma:<br>- Shape: 1<br>- Scale: 200,000<br>- Offset: 0<br>Uniform:<br>- Lower: 36913,000<br>- Upper: 786,913<br>Initial: 100,000 | Point estimate: 56,8087 |
| YG 26.1 | Gamma:<br>- Shape: 1<br>- Scale: 200,000<br>- Offset: 0<br>Uniform:<br>- Lower: 36913,000<br>- Upper: 786,913<br>Initial: 100,000 | Point estimate: 116,087 |
| UAMES 9705 | Gamma:<br>- Shape: 1<br>- Scale: 200,000<br>- Offset: 0<br>Uniform:<br>- Lower: 36913,000<br>- Upper: 786,913<br>Initial: 100,000 | Point estimate: 135,087 |
| UAMES 7663 | Gamma:<br>- Shape: 1<br>- Scale: 200,000<br>- Offset: 0<br>Uniform:<br>- Lower: 36913,000<br>- Upper: 786,913<br>Initial: 100,000 | Point estimate: 124,087 |
| YG 43.2 | Gamma:<br>- Shape: 1<br>- Scale: 200,000<br>- Offset: 0<br>Uniform:<br>- Lower: 36913,000<br>- Upper: 786,913<br>Initial: 100,000 | Point estimate: 135,087 |
| UM58075 | Point estimate: 13,881 | Point estimate: 13,881 |
| UM57705 | Point estimate: 15,334 | Point estimate: 15,334 |

| | | |
|---|---|---|
| RAM P97.7.1 | Gamma:<br>- Shape: 1<br>- Scale: 200,000<br>- Offset: 0<br>Uniform:<br>- Lower: 36913,000<br>- Upper: 786,913<br>Initial: 100,000 | Point estimate: 183,087 |
| YG 50.1 | Gamma:<br>- Shape: 1<br>- Scale: 200,000<br>- Offset: 0<br>Uniform:<br>- Lower: 36913,000<br>- Upper: 786,913<br>Initial: 100,000 | Point estimate: 102,951 |
| UAMES 30198 | Gamma:<br>- Shape: 1<br>- Scale: 200,000<br>- Offset: 0<br>Uniform:<br>- Lower: 36913,000<br>- Upper: 786,913<br>Initial: 100,000 | Point estimate: 132,087 |
| UAMES 30197 | Gamma:<br>- Shape: 1<br>- Scale: 200,000<br>- Offset: 0<br>Uniform:<br>- Lower: 36913,000<br>- Upper: 786,913<br>Initial: 100,000 | Point estimate: 541,087 |
| UAMES 30199 | Gamma:<br>- Shape: 1<br>- Scale: 200,000<br>- Offset: 0<br>Uniform:<br>- Lower: 36913,000<br>- Upper: 786,913<br>Initial: 100,000 | Point estimate: 135,087 |
| RAM P94.16.1B | Gamma:<br>- Shape: 1<br>- Scale: 200,000<br>- Offset: 0<br>Uniform:<br>- Lower: 36913,000<br>- Upper: 786,913<br>Initial: 100,000 | Point estimate: 216,087 |

| | | |
|---|---|---|
| RAM P94.5.7 | Gamma:<br>- Shape: 1<br>- Scale: 200,000<br>- Offset: 0<br>Uniform:<br>- Lower: 36913,000<br>- Upper: 786,913<br>Initial: 100,000 | Point estimate: 469,087 |
| UAMES 12060 | Gamma:<br>- Shape: 1<br>- Scale: 200,000<br>- Offset: 0<br>Uniform:<br>- Lower: 36913,000<br>- Upper: 786,913<br>Initial: 100,000 | Point estimate: 124,087 |
| UAMES 34125 | Gamma:<br>- Shape: 1<br>- Scale: 200,000<br>- Offset: 0<br>Uniform:<br>- Lower: 36913,000<br>- Upper: 786,913<br>Initial: 100,000 | Point estimate: 134,087 |
| INSM 71.3.261 | Point estimate: 13,326 | Point estimate: 13,326 |
| F:AM 103291 | Gamma:<br>- Shape: 1<br>- Scale: 200,000<br>- Offset: 0<br>Uniform:<br>- Lower: 36913,000<br>- Upper: 786,913<br>Initial: 100,000 | Point estimate: 115,087 |
| P 14591 | Point estimate: 13,229 | Point estimate: 13,229 |
| ETMNH 19335 | Point estimate: 26,708 | Point estimate: 26,708 |
| ETMNH 19334 | Point estimate: 24,872 | Point estimate: 24,872 |
| ISM 65BS68 | Point estimate: 13,493 | Point estimate: 13,493 |
| UWZM 19581 | Point estimate: 13,099 | Point estimate: 13,099 |
| UWZM 19580 | Point estimate: 13,087 | Point estimate: 13,087 |
| DP1296 | Point estimate: 34,329 | Point estimate: 34,329 |
| IMNH 50003/16643 | Point estimate: 49,636 | Point estimate: 49,636 |
| IMNH 52002/25939 | Gamma:<br>- Shape: 1<br>- Scale: 200,000<br>- Offset: 0<br>Uniform:<br>- Lower: 0<br>- Upper: 786,913<br>Initial: 20,000 | Point estimate: 64,414 |

| | | |
|---|---|---|
| NSM979GF153.001 | Gamma:<br>- Shape: 1<br>- Scale: 200,000<br>- Offset: 0<br>Uniform:<br>- Lower: 36,913<br>- Upper: 786,913<br>Initial: 100,000 | Gamma:<br>- Shape: 1<br>- Scale: 200,000<br>- Offset: 0<br>Uniform:<br>- Lower: 36,913<br>- Upper: 786,913<br>Initial: 100,000 |
| NSM989GF089.002 | Gamma:<br>- Shape: 1<br>- Scale: 200,000<br>- Offset: 0<br>Uniform:<br>- Lower: 36,913<br>- Upper: 786,913<br>Initial: 100,000 | Gamma:<br>- Shape: 1<br>- Scale: 200,000<br>- Offset: 0<br>Uniform:<br>- Lower: 36,913<br>- Upper: 786,913<br>Initial: 100,000 |
| NSM019GF009.002 | Point estimate: 74,900 | Point estimate: 74,900 |
| NSM015GF007 | Gamma:<br>- Shape: 1<br>- Scale: 200,000<br>- Offset: 0<br>Uniform:<br>- Lower: 36,913<br>- Upper: 786,913<br>Initial: 100,000 | Gamma:<br>- Shape: 1<br>- Scale: 200,000<br>- Offset: 0<br>Uniform:<br>- Lower: 36,913<br>- Upper: 786,913<br>Initial: 100,000 |
| NSM021GF005.001 | Gamma:<br>- Shape: 1<br>- Scale: 200,000<br>- Offset: 0<br>Uniform:<br>- Lower: 0<br>- Upper: 786,913<br>Initial: 20,000 | Gamma:<br>- Shape: 1<br>- Scale: 200,000<br>- Offset: 0<br>Uniform:<br>- Lower: 36,913<br>- Upper: 786,913<br>Initial: 100,000 |
| Renison | Gamma:<br>- Shape: 1<br>- Scale: 200,000<br>- Offset: 0<br>Uniform:<br>- Lower: 36,913<br>- Upper: 786,913<br>Initial: 100,000 | Gamma:<br>- Shape: 1<br>- Scale: 200,000<br>- Offset: 0<br>Uniform:<br>- Lower: 36,913<br>- Upper: 786,913<br>Initial: 100,000 |
| Tualatin | Point estimate: 13,360 | Point estimate: 13,360 |
| NC_035800 | Point estimate: 13,401 | Point estimate: 13,401 |
| EF_632344 | Gamma:<br>- Shape: 1<br>- Scale: 200,000<br>- Offset: 0<br>Uniform:<br>- Lower: 36,913<br>- Upper: 786,913<br>Initial: 100,000 | Point estimate: 135,087 |

**Supplementary Table 3.** Data table of all parameters from dating analyses when non-finite or undated mastodons are fitted with broad gamma distributions. Values are after burn-in has been removed and all three independent chains combined. All age values are uncorrected and relative to the youngest specimen (UWZM 19580 – 13087 yBP).

| Summary Statistic | joint | prior | likelihood | treeModel.rootHeight | age(root) | treeLength | tmrca(Unknowns) | age(Unknowns) |
|---|---|---|---|---|---|---|---|---|
| mean | -29926.8 | -1373.67 | -28553.2 | 2.71E+06 | 2.70E+06 | 1.21E+07 | 2.71E+06 | 2.70E+06 |
| stderr of mean | 0.1921 | 0.1874 | 0.0238 | 7171.351 | 7171.351 | 32197.43 | 7171.302 | 7171.302 |
| stdev | 13.34 | 11.2781 | 6.023 | 4.95E+05 | 4.95E+05 | 2.17E+06 | 4.95E+05 | 4.95E+05 |
| variance | 177.9551 | 127.1945 | 36.2769 | 2.45E+11 | 2.45E+11 | 4.70E+12 | 2.45E+11 | 2.45E+11 |
| median | -29927.5 | -1374.63 | -28552.9 | 2.70E+06 | 2.69E+06 | 1.21E+07 | 2.70E+06 | 2.69E+06 |
| value range | [-29979.4922, -29863.361] | [-1412.8679, -1313.8135] | [-28588.3239, -28530.8601] | [9.5062E5, 5.0187E6] | [9.3753E5, 5.0056E6] | [4.2276E6, 2.1898E7] | [9.5062E5, 5.0187E6] | [9.3753E5, 5.0056E6] |
| geometric mean | n/a | n/a | n/a | 2.66E+06 | 2.65E+06 | 1.19E+07 | 2.66E+06 | 2.65E+06 |
| 95% HPD interval | [-29952.0185, -29899.6233] | [-1394.7883, -1350.8159] | [-28565.3001, -28541.7653] | [1.7236E6, 3.6694E6] | [1.7105E6, 3.6563E6] | [7.8522E6, 1.6363E7] | [1.7235E6, 3.6694E6] | [1.7104E6, 3.6563E6] |
| auto-correlation time (ACT) | 2.80E+05 | 3.73E+05 | 21154.15 | 2.84E+05 | 2.84E+05 | 2.98E+05 | 2.84E+05 | 2.84E+05 |
| effective sample size (ESS) | 4822.1 | 3621.3 | 63818.7 | 4759.9 | 4759.9 | 4529.4 | 4760.1 | 4760.1 |
| number of samples | 1.35E+05 | 1.35E+05 | 1.35E+05 | 1.35E+05 | 1.35E+05 | 1.35E+05 | 1.35E+05 | 1.35E+05 |

| constant.popSize | kappa | alpha | clock.rate | meanRate | age(52002_25939_consensus) | age(AMNH_988_consensus_sequence) | age(CMN_11697_consensus_sequence) |
|---|---|---|---|---|---|---|---|
| 7.42E+05 | 49.0506 | 0.0536 | 5.55E-09 | 5.55E-09 | 49992.91 | 19049.72 | 1.17E+05 |
| 1727.995 | 0.0212 | 9.73E-05 | 1.73E-11 | 1.73E-11 | 74.0448 | 63.9726 | 418.4622 |
| 1.49E+05 | 7.8052 | 0.0348 | 1.11E-09 | 1.11E-09 | 16924.19 | 16538.04 | 37947.35 |
| 2.23E+10 | 60.9204 | 1.21E-03 | 1.23E-18 | 1.23E-18 | 2.86E+08 | 2.74E+08 | 1.44E+09 |
| 7.51E+05 | 48.2742 | 0.0486 | 5.36E-09 | 5.36E-09 | 49041.94 | 14634.87 | 1.14E+05 |
| [1.8661E5, 9.9998E5] | [24.8498, 103.4363] | [2.7904E-3, 0.2885] | [2.9409E-9, 1.5273E-8] | [2.9409E-9, 1.5273E-8] | [25.4034, 1.6661E5] | [0.0488, 1.4825E5] | [36915.7777, 3.0726E5] |
| 7.25E+05 | 48.4503 | 0.0405 | 5.45E-09 | 5.45E-09 | 46452.21 | 11585.14 | 1.10E+05 |
| [4.8249E5, 9.9998E5] | [34.7681, 64.6357] | [2.7904E-3, 0.117] | [3.7382E-9, 7.8112E-9] | [3.7382E-9, 7.8112E-9] | [15997.6182, 85930.5745] | [0.0488, 51864.3303] | [46761.9806, 1.9039E5] |
| 1.81E+05 | 10000 | 10519.42 | 3.29E+05 | 3.29E+05 | 25841.62 | 20200.69 | 1.64E+05 |
| 7472.8 | 135003 | 128336.9 | 4104.3 | 4104.3 | 52242.5 | 66830.9 | 8223.3 |
| 1.35E+05 | 1.35E+05 | 1.35E+05 | 1.35E+05 | 1.35E+05 | 1.35E+05 | 1.35E+05 | 1.35E+05 |

**Supplementary Table 3.** Continued

| age(FAM_103291_consensus_sequence) | age(IK-01-277_consensus_sequence) | age(IK-01-321_consensus_sequence) | age(IK-98-963_consensus_sequence) | age(IK-99-328_consensus_sequence) | age(IK05-3.5_consensus_sequence) | age(IK08-127_consensus_sequence) | age(IK10-106_consensus_sequence) |
|---:|---:|---:|---:|---:|---:|---:|---:|
| 1.00E+05 | 1.08E+05 | 1.17E+05 | 1.08E+05 | 4.85E+05 | 4.63E+05 | 1.14E+05 | 1.08E+05 |
| 330.751 | 361.8876 | 417.4096 | 359.7369 | 3362.15 | 3320.174 | 401.7259 | 362.4058 |
| 36652.3 | 37609.18 | 37885.3 | 37655.27 | 1.20E+05 | 1.20E+05 | 38111.22 | 37471.71 |
| 1.34E+09 | 1.41E+09 | 1.44E+09 | 1.42E+09 | 1.44E+10 | 1.43E+10 | 1.45E+09 | 1.40E+09 |
| 96236.69 | 1.05E+05 | 1.14E+05 | 1.05E+05 | 4.85E+05 | 4.62E+05 | 1.11E+05 | 1.05E+05 |
| [36913.968, 2.8664E5] | [36914.3727, 3.265E5] | [36921.6944, 3.2966E5] | [36919.5399, 3.2869E5] | [41949.8911, 7.8686E5] | [38073.49, 7.8679E5] | [36917.9577, 3.4039E5] | [36919.1385, 3.2454E5] |
| 93423.16 | 1.02E+05 | 1.11E+05 | 1.02E+05 | 4.69E+05 | 4.46E+05 | 1.08E+05 | 1.02E+05 |
| [36913.968, 1.6596E5] | [37136.6481, 1.7529E5] | [46912.9431, 1.9013E5] | [37093.4711, 1.7579E5] | [2.5251E5, 7.2013E5] | [2.3187E5, 6.989E5] | [42975.0491, 1.8608E5] | [38027.7498, 1.7601E5] |
| 1.10E+05 | 1.25E+05 | 1.64E+05 | 1.23E+05 | 1.06E+06 | 1.04E+06 | 1.50E+05 | 1.26E+05 |
| 12280 | 10800.3 | 8237.8 | 10956.7 | 1270.3 | 1297.3 | 9000 | 10690.9 |
| 1.35E+05 | 1.35E+05 | 1.35E+05 | 1.35E+05 | 1.35E+05 | 1.35E+05 | 1.35E+05 | 1.35E+05 |

| age(KIG12-15_consensus_sequence) | age(MAY12-69_consensus_sequence) | age(MAY12-70_consensus_sequence) | age(NC_009574) | age(NSM015GF007_curated) | age(NSM021GF005_001_curated) | age(NSM979GF153_001_curated) | age(NSM989GF089_002_curated) |
|---:|---:|---:|---:|---:|---:|---:|---:|
| 1.17E+05 | 1.17E+05 | 1.17E+05 | 1.17E+05 | 6.27E+05 | 29350.84 | 93680.03 | 2.96E+05 |
| 425.0115 | 427.2913 | 428.3573 | 430.3227 | 1455.251 | 107.216 | 138.7947 | 718.9493 |
| 37847.16 | 38073.29 | 38223.5 | 37717.7 | 98365.09 | 21582.65 | 34094.2 | 76169.33 |
| 1.43E+09 | 1.45E+09 | 1.46E+09 | 1.42E+09 | 9.68E+09 | 4.66E+08 | 1.16E+09 | 5.80E+09 |
| 1.14E+05 | 1.14E+05 | 1.14E+05 | 1.14E+05 | 6.37E+05 | 25386.04 | 89684.74 | 2.94E+05 |
| [36913.1651, 3.2104E5] | [36918.6631, 3.4223E5] | [36913.5885, 3.292E5] | [36914.3002, 3.1202E5] | [1.6025E5, 7.8691E5] | [0.2691, 1.808E5] | [36913.1586, 3.1646E5] | [39090.803, 6.6946E5] |
| 1.11E+05 | 1.11E+05 | 1.11E+05 | 1.11E+05 | 6.19E+05 | 19802.75 | 87658.72 | 2.86E+05 |
| [46423.2696, 1.8975E5] | [47114.7919, 1.9116E5] | [46812.765, 1.9109E5] | [47204.8938, 1.8996E5] | [4.5112E5, 7.869E5] | [0.2691, 70506.7074] | [36913.1586, 1.559E5] | [1.5124E5, 4.4749E5] |
| 1.70E+05 | 1.70E+05 | 1.70E+05 | 1.76E+05 | 2.95E+05 | 33316.26 | 22373.38 | 1.20E+05 |
| 7929.8 | 7939.4 | 7962.4 | 7682.4 | 4568.8 | 40521.7 | 60340.9 | 11224.3 |
| 1.35E+05 | 1.35E+05 | 1.35E+05 | 1.35E+05 | 1.35E+05 | 1.35E+05 | 1.35E+05 | 1.35E+05 |

**Supplementary Table 3.** Continued

| age(RAM_P94.16.1B_consensus_sequence) | age(RAM_P94.5.7_consensus_sequence) | age(RAM_P97.7.1_consensus_sequence) | age(Renison_curated) | age(UAMES_7663_consensus_sequence) | age(UM13909_consensus_sequence) | age(YG26.1_consensus_sequence) | age(YG43.2_consensus_sequence) |
|---|---|---|---|---|---|---|---|
| 1.78E+05 | 3.46E+05 | 3.63E+05 | 2.85E+05 | 1.07E+05 | 28619.32 | 1.01E+05 | 1.17E+05 |
| 468.1034 | 854.2658 | 1076.599 | 695.268 | 362.6207 | 94.6552 | 315.9437 | 424.0829 |
| 37243.43 | 72060.33 | 1.22E+05 | 81034.24 | 37623.14 | 20354.53 | 36513.87 | 38021.09 |
| 1.39E+09 | 5.19E+09 | 1.48E+10 | 6.57E+09 | 1.42E+09 | 4.14E+08 | 1.33E+09 | 1.45E+09 |
| 1.77E+05 | 3.44E+05 | 3.52E+05 | 2.83E+05 | 1.04E+05 | 25158.31 | 96919.85 | 1.14E+05 |
| [41316.404, 3.868E5] | [89599.9494, 7.0459E5] | [37649.1986, 7.8672E5] | [37020.1129, 6.8882E5] | [36920.9736, 3.1711E5] | [0.0673, 1.554E5] | [36915.0768, 3.4878E5] | [36921.396, 3.2035E5] |
| 1.74E+05 | 3.39E+05 | 3.42E+05 | 2.73E+05 | 1.01E+05 | 19513.56 | 94013.46 | 1.11E+05 |
| [1.0665E5, 2.5191E5] | [2.0917E5, 4.8949E5] | [1.4568E5, 6.0767E5] | [1.3139E5, 4.4627E5] | [37000.1753, 1.747E5] | [0.0673, 66622.195] | [36924.1237, 1.6662E5] | [46475.121, 1.9003E5] |
| 2.13E+05 | 1.90E+05 | 1.06E+05 | 99383.44 | 1.25E+05 | 29195.32 | 1.01E+05 | 1.68E+05 |
| 6330.1 | 7115.5 | 12780.2 | 13584.1 | 10764.7 | 46241.3 | 13356.5 | 8037.9 |
| 1.35E+05 | 1.35E+05 | 1.35E+05 | 1.35E+05 | 1.35E+05 | 1.35E+05 | 1.35E+05 | 1.35E+05 |

| age(YG50.1_consensus_sequence) | treeLikelihood | branchRates | coalescent |
|---|---|---|---|
| 90115.02 | -28553.2 | 0 | -623.743 |
| 274.6682 | 0.0238 | 0 | 0.1306 |
| 34654.44 | 6.023 | 0 | 8.9327 |
| 1.20E+09 | 36.2769 | 0 | 79.7939 |
| 85285.42 | -28552.9 | 0 | -624.495 |
| [36914.7502, 3.1148E5] | [-28588.3239, -28530.8601] | [0, 0] | [-656.4907, -574.991] |
| 83711.26 | n/a | n/a | n/a |
| [36916.4285, 1.5421E5] | [-28565.3001, -28541.7653] | n/a | [-640.9249, -605.9573] |
| 84809.69 | 21154.15 | n/a | 2.89E+05 |
| 15918.3 | 63818.7 | n/a | 4676.8 |
| 1.35E+05 | 1.35E+05 | 1.35E+05 | 1.35E+05 |

**Supplementary Figure 10.** MCC tree with node age estimates when mastodons with known dates are used to calibrate the phylogeny.

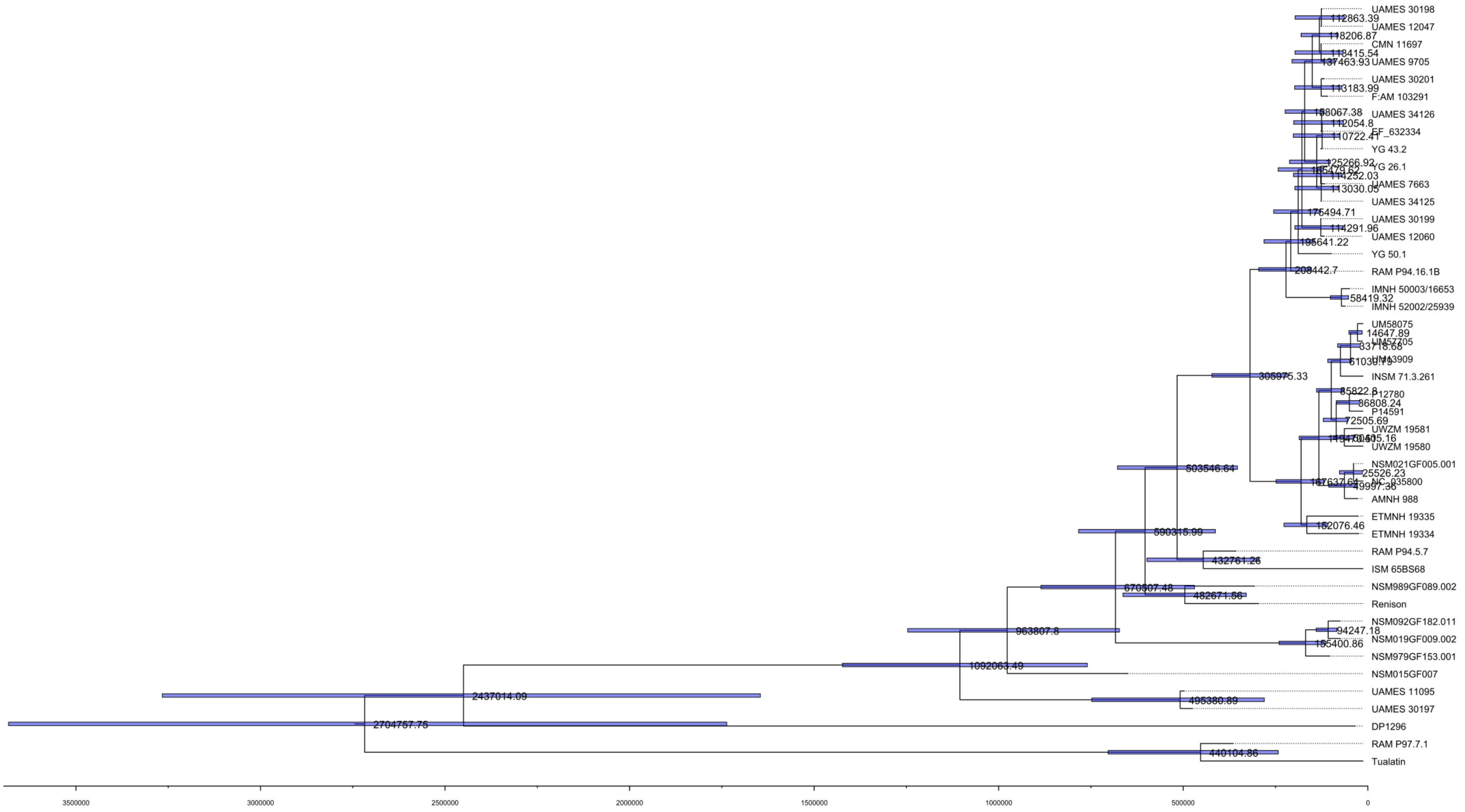

**Supplementary Table 4.** Data table of all parameters from dating analyses when non-finite or undated mastodons are fitted with broad gamma distributions. Values are after burn-in has been removed and all three independent chains combined. All age values are uncorrected and relative to the youngest specimen (UWZM 19580 – 13087 yBP).

| Summary Statistic | joint | prior | likelihood | treeModel.rootHeight | age(root) | treeLength | tmrca(Unknowns) |
|---|---|---|---|---|---|---|---|
| mean | -29312.3 | -765.062 | -28547.2 | 2.64E+06 | 2.62E+06 | 1.21E+07 | 1.02E+06 |
| stderr of mean | 0.0242 | 0.0193 | 0.014 | 1042.469 | 1042.469 | 4649.492 | 276.7829 |
| stdev | 7.1027 | 4.5959 | 5.127 | 2.60E+05 | 2.60E+05 | 1.11E+06 | 74759.94 |
| variance | 50.449 | 21.122 | 26.2861 | 6.74E+10 | 6.74E+10 | 1.24E+12 | 5.59E+09 |
| median | -29311.9 | -764.972 | -28547 | 2.62E+06 | 2.60E+06 | 1.20E+07 | 1.02E+06 |
| value range | [-29350.399, -29288.1291] | [-786.7562, -744.2268] | [-28576.4156, -28528.8572] | [1.7716E6, 4.0185E6] | [1.7585E6, 4.0054E6] | [8.2946E6, 1.7958E7] | [7.5261E5, 1.3995E6] |
| geometric mean | n/a | n/a | n/a | 2.62E+06 | 2.61E+06 | 1.20E+07 | 1.02E+06 |
| 95% HPD interval | [-29326.4946, -29298.8458] | [-773.8918, -755.8664] | [-28557.436, -28537.5193] | [2.1432E6, 3.1441E6] | [2.1301E6, 3.131E6] | [9.9664E6, 1.4268E7] | [8.7952E5, 1.1721E6] |
| auto-correlation time (ACT) | 15721.83 | 23690.32 | 10102.25 | 21764.72 | 21764.72 | 23611.1 | 18504.92 |
| effective sample size (ESS) | 85869.8 | 56986.6 | 133636.6 | 62028.4 | 62028.4 | 57177.8 | 72955.2 |
| number of samples | 1.35E+05 | 1.35E+05 | 1.35E+05 | 1.35E+05 | 1.35E+05 | 1.35E+05 | 1.35E+05 |

| age(Unknowns) | constant.popSize | kappa | alpha | clock.rate | meanRate | age(NSM015GF007_curated) |
|---|---|---|---|---|---|---|
| 1.01E+06 | 7.54E+05 | 48.9358 | 0.0556 | 5.43E-09 | 5.43E-09 | 6.86E+05 |
| 276.7829 | 377.4774 | 0.0212 | 9.89E-05 | 2.13E-12 | 2.13E-12 | 190.5143 |
| 74759.94 | 1.18E+05 | 7.785 | 0.0362 | 5.25E-10 | 5.25E-10 | 66978.99 |
| 5.59E+09 | 1.40E+10 | 60.606 | 1.31E-03 | 2.76E-19 | 2.76E-19 | 4.49E+09 |
| 1.00E+06 | 7.52E+05 | 48.1543 | 0.0508 | 5.41E-09 | 5.41E-09 | 6.95E+05 |
| [7.3952E5, 1.3864E6] | [3.7079E5, 1E6] | [25.4403, 109.4836] | [2.7911E-3, 0.2798] | [3.4224E-9, 7.956E-9] | [3.4224E-9, 7.956E-9] | [2.6449E5, 7.8691E5] |
| 1.00E+06 | 7.45E+05 | 48.3368 | 0.0419 | 5.40E-09 | 5.40E-09 | 6.82E+05 |
| [8.6643E5, 1.159E6] | [5.5377E5, 9.9076E5] | [34.6433, 64.3443] | [2.7911E-3, 0.1218] | [4.4217E-9, 6.4704E-9] | [4.4217E-9, 6.4704E-9] | [5.6221E5, 7.8691E5] |
| 18504.92 | 13721.67 | 10028.38 | 10098.57 | 22088.34 | 22088.34 | 10922.56 |
| 72955.2 | 98386.7 | 134621 | 133685.3 | 61119.6 | 61119.6 | 123600.1 |
| 1.35E+05 | 1.35E+05 | 1.35E+05 | 1.35E+05 | 1.35E+05 | 1.35E+05 | 1.35E+05 |

**Supplementary Table 4.** Continued

| age(NSM021GF005_001_curated) | age(NSM979GF153_001_curated) | age(NSM989GF089_002_curated) | age(Renison_curated) | treeLikelihood | branchRates | coalescent |
|---|---|---|---|---|---|---|
| 28449.27 | 99224.33 | 3.48E+05 | 3.39E+05 | -28547.2 | 0 | -624.276 |
| 61.2714 | 99.6291 | 181.3403 | 199.8772 | 0.014 | 0 | 0.0187 |
| 19902.16 | 36453.63 | 65657.97 | 72722.91 | 5.127 | 0 | 4.5261 |
| 3.96E+08 | 1.33E+09 | 4.31E+09 | 5.29E+09 | 26.2861 | 0 | 20.4859 |
| 25491.35 | 95549.06 | 3.50E+05 | 3.41E+05 | -28547 | 0 | -624.188 |
| [0.5639, 1.5898E5] | [36913.5095, 3.1814E5] | [38376.1818, 6.518E5] | [37610.0683, 6.3694E5] | [-28576.4156, -28528.8572] | [0, 0] | [-644.615, -604.5978] |
| 19662.59 | 92531.27 | 3.42E+05 | 3.30E+05 | n/a | n/a | n/a |
| [0.5639, 65775.7335] | [36913.5095, 1.6409E5] | [2.1779E5, 4.7576E5] | [1.9508E5, 4.8037E5] | [-28557.436, -28537.5193] | n/a | [-633.2395, -615.5149] |
| 12795.63 | 10084.12 | 10298.18 | 10198.36 | 10102.25 | n/a | 23031.18 |
| 105507.1 | 133876.9 | 131094 | 132377.1 | 133636.6 | n/a | 58617.5 |
| 1.35E+05 | 1.35E+05 | 1.35E+05 | 1.35E+05 | 1.35E+05 | 1.35E+05 | 1.35E+05 |

**Supplementary Figure 11.** MCC tree with node posterior probabilities shown when all previous sequenced mastodons are fit with point estimates equal to the median calibrated radiocarbon age or the median estimated age in Karpinski et al. (2023).

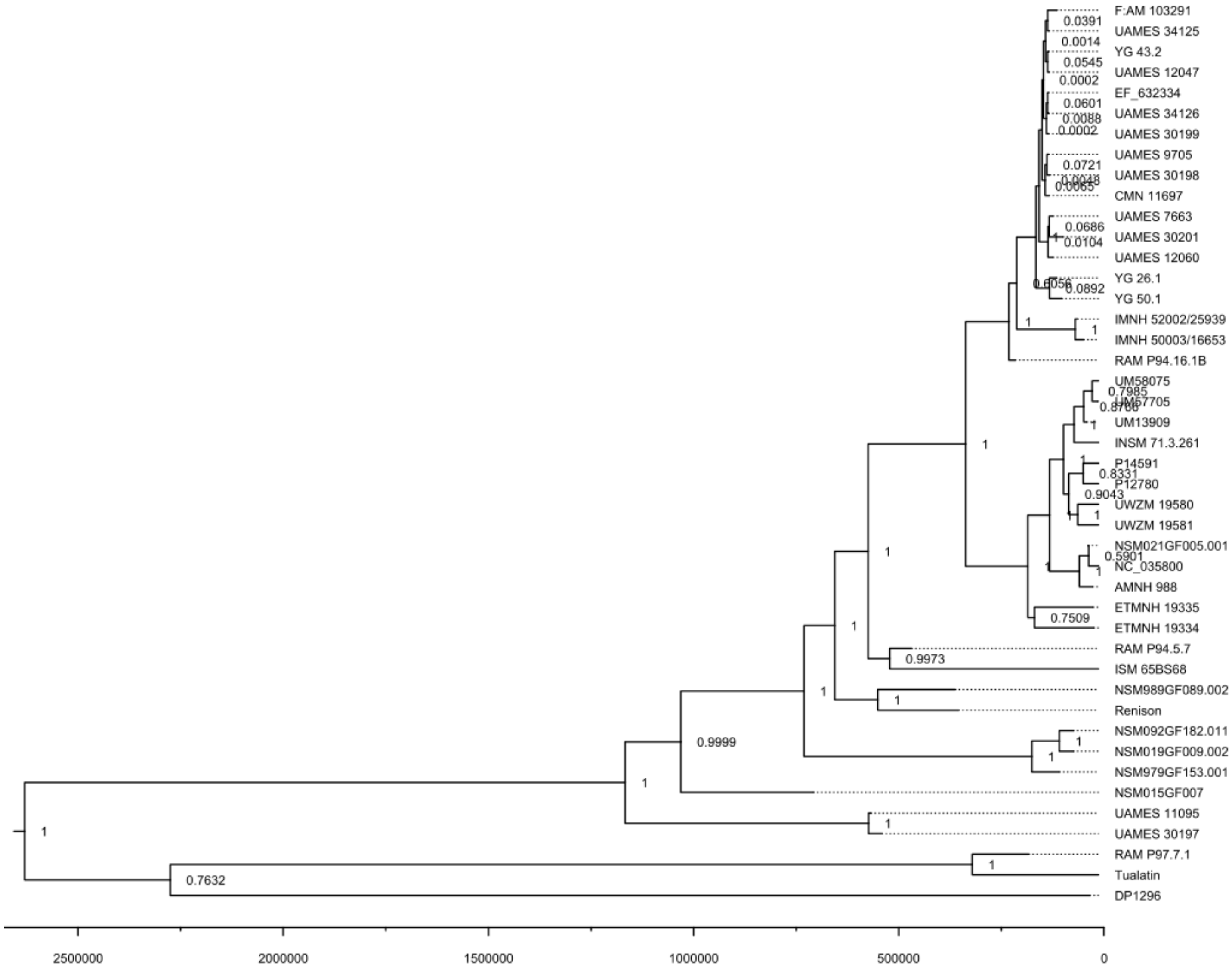

**Supplementary Figure 12.** MCC tree with node age estimates shown when all previous sequenced mastodons are fit with point estimates equal to the median calibrated radiocarbon age or the median estimated age in Karpinski et al. (2023).

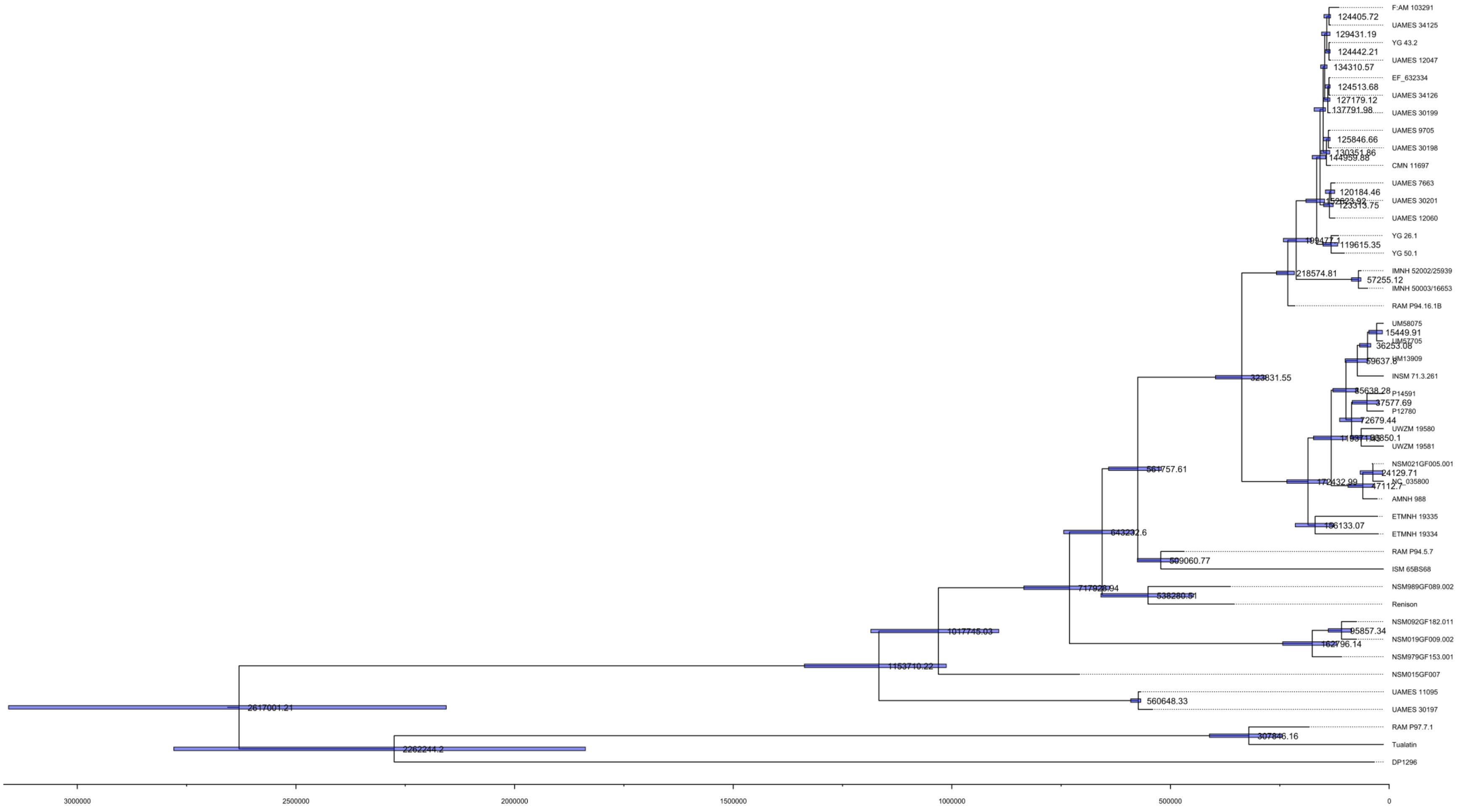

**Supplementary Figure 13.** Violin plot with the posterior probability distributions on ages of the samples within this study when all previous sequenced mastodons are fit with point estimates equal to the median calibrated radiocarbon age or the media estimated age in Karpinski et al. (2023). Ages on the y-axis are relative to the youngest sample in the analysis (UWZM 19580 – 13087 yBP)

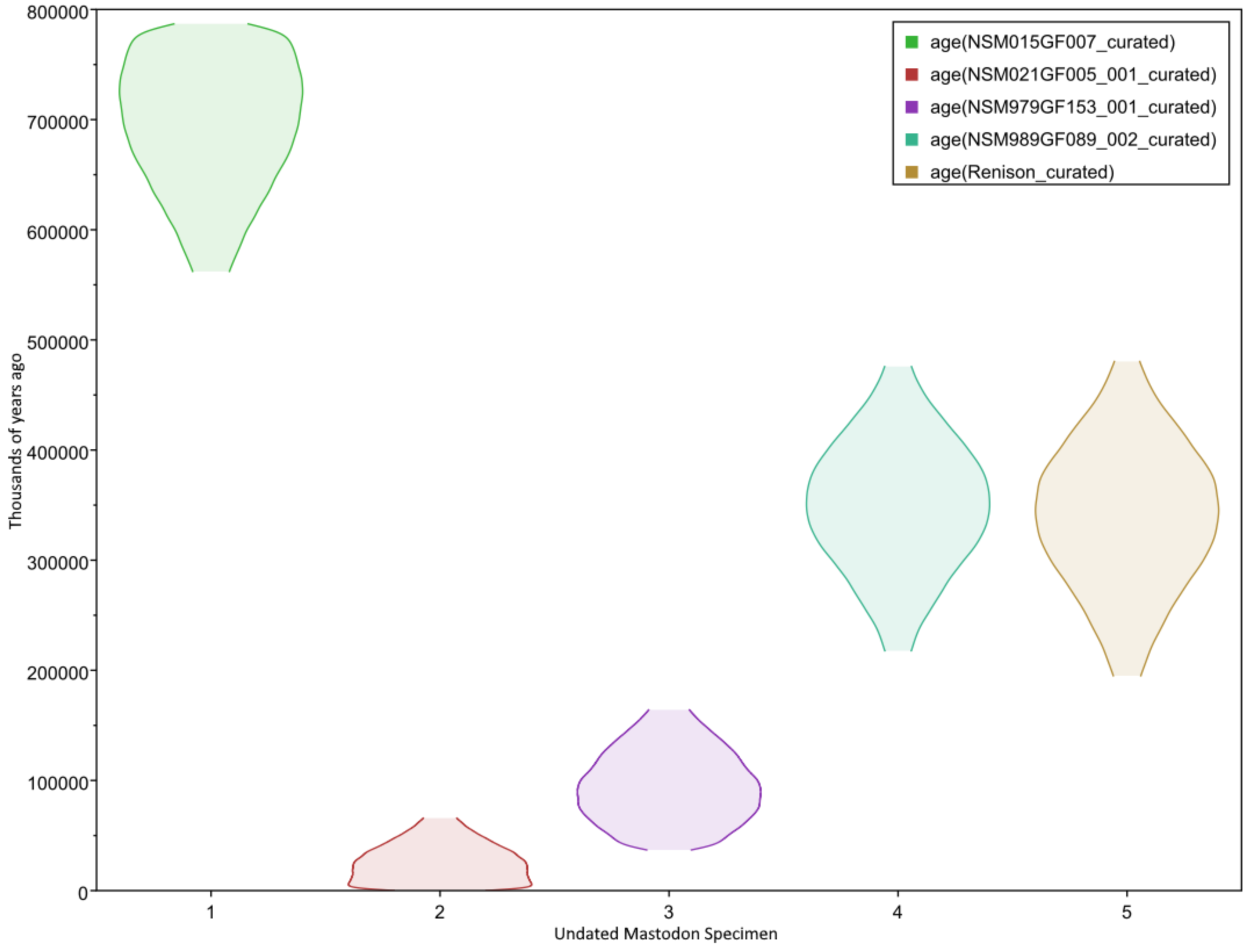

**Supplementary Table 5.** Details of new indirect radiocarbon dates reported in this study.

| University of California # | Université Laval # | Cust. ID (sample type) | Pre-Treatment | $F^{14}C$ | ± | $D^{14}C$ (‰) | ± | $^{14}C$ age (BP) | ± |
|---|---|---|---|---|---|---|---|---|---|
| UCIAMS-157040 | ULA-5275 | LN-X1 (wood) | HCl-NaOH-HCl | -0.0006 | -0.0007 | -1000.6 | -0.7 | >52800 | |
| UCIAMS-157044 | ULA-5276 | LN-S2 (wood) | HCl-NaOH-HCl | 0.0005 | 0.0007 | -999.5 | 0.7 | >50300 | |

**Supplementary Table 6.** Details of new direct radiocarbon dates reported in this study.

| University of California # | Université Laval # | Cust. ID (sample type) | Pre-Treatment | $F^{14}C$ | ± | $D^{14}C$ (‰) | ± | $^{14}C$ age (BP) | ± | $\delta^{13}C$ (‰) | % N | % C | C/N (wt%/ wt%) | % of bone transformed into collagen |
|---|---|---|---|---|---|---|---|---|---|---|---|---|---|---|
| UCIAMS-157082 | ULA-5288 | LN-B1 (collagen from bone) | HCl-NaOH-HCl | 0.0062 | 0.0017 | -993.8 | 1.7 | 40900 | 2300 | -20.9 | 15.5 | 45.3 | 2.91 | 9.3 |
